# Sex-dependent control of renal tubular homeostasis and stress tolerance by KDM6A

**DOI:** 10.64898/2026.07.22.740137

**Authors:** Lisa Y. Q. Hong, Sri Nagarjun Batchu, Duc Tin Tran, Madiha Zahra Syeda, Suzanne L. Advani, Youan Liu, Alain Pacis, Evgeniy V. Petrotchenko, Christoph H. Borchers, Darren A. Yuen, Andrew Advani

**Author notes:** Address for correspondence: Dr. Andrew Advani, St. Michael’s Hospital, 6-151 61 Queen Street East Toronto, Ontario, Canada, M5C 2T2.

## Abstract

Biological sex is an important determinant of kidney disease susceptibility and outcomes. The epigenetic modifier KDM6A is an X chromosome-expressed lysine demethylase and molecular scaffold that escapes X chromosome inactivation. Here, we compared the effects of deletion of KDM6A from kidney tubule epithelial cells in female and male mice (KDM6A^TubKO^). Knockout of KDM6A from tubule cells aggravated kidney fibrosis caused by unilateral ureteral obstruction (UUO) in female mice, whereas male mice were unaffected by KDM6A absence. Unexpectedly, female (but not male) KDM6A^TubKO^ mice developed spontaneous glucosuria that, when stressed by ligation of one ureter, presented as polyuria and a Fanconi renotubular syndrome-like picture affecting the unobstructed kidney. Absence of KDM6A from tubule epithelial cells of female mice caused mitochondrial circularization, tubule cell vacuolization with focal atrophy and lymphoid infiltration, and diminished sodium/glucose cotransporter 2 (SGLT2). Spatial transcriptomics and untargeted metabolomics revealed that knockout of KDM6A in female mice caused a shift in gene programs and metabolic pathways indicative of tubule cell metabolic dysfunction. In male mice, transcripts of the Y chromosome-expressed gametolog of *Kdm6a*, *Uty* were present in tubule epithelial cells at levels comparable to *Kdm6a* and they were upregulated with UUO. In summary, KDM6A is essential for normal tubule epithelial cell homeostasis in females but not in males. KDM6A and UTY are a dynamically regulated X-Y gene pair with at least partial compensatory overlap in function necessary for the preservation of kidney health and stress tolerance.

**Graphical abstract.:** 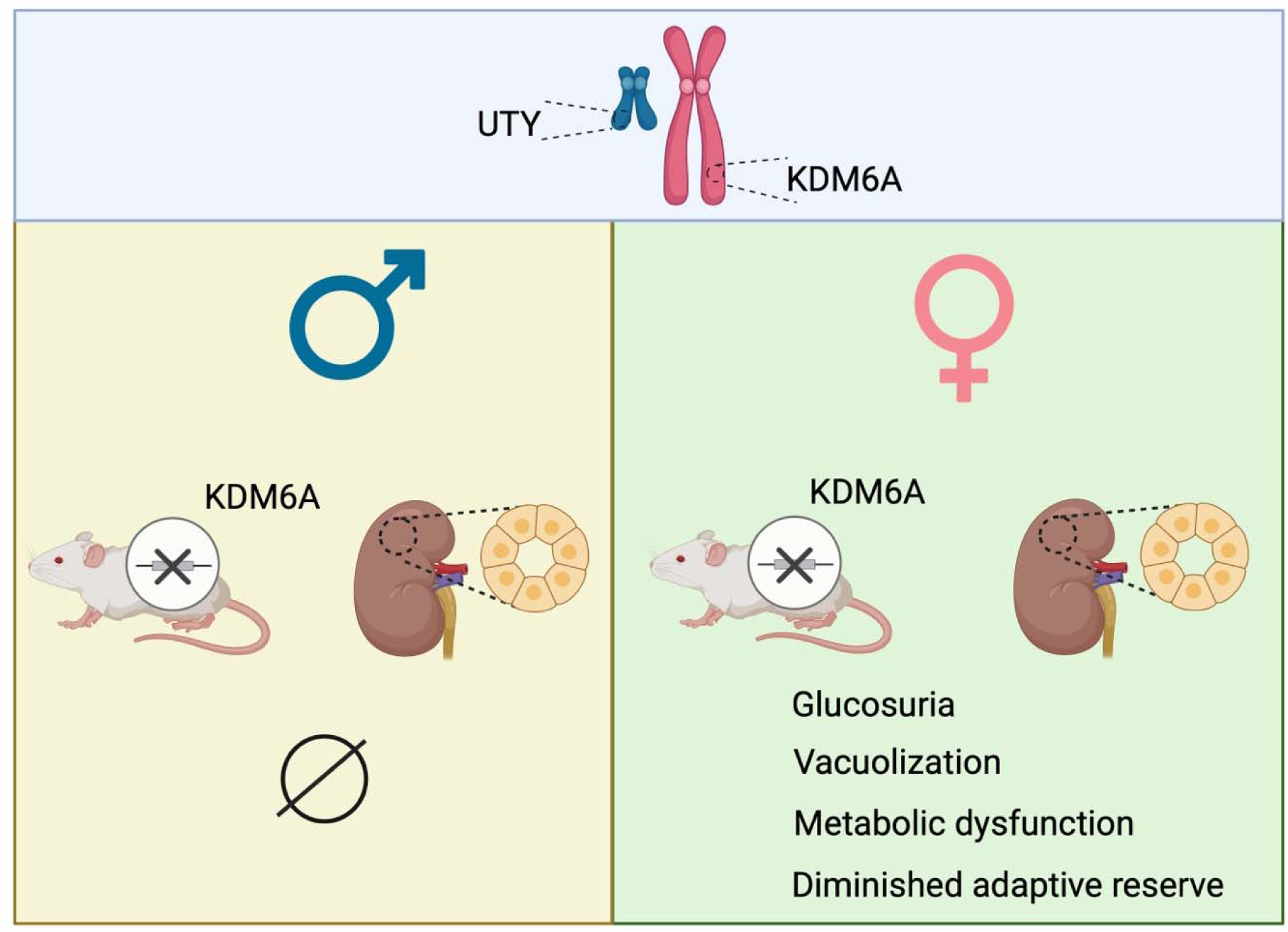

---

Although it was historically often overlooked, there is now growing recognition that sex has an important influence on both the development and outcomes of kidney disease. For instance, chronic kidney disease (CKD) is more prevalent in adult women than adult men, yet CKD outcomes are worse in adult men than women.^1^ Conversely, in pediatric populations, kidney disease is more common in boys than girls, largely because of an increased prevalence of congenital abnormalities of the kidney and urinary tract (CAKUT) in boys.^2^ Sex differences in kidney disease may arise from a complex interplay of social factors like access to care,^1^ hormonal factors like estrogens and androgens ^3^ and genetic factors like genes expressed by the sex chromosomes. Lysine-specific demethylase 6A (KDM6A) is expressed on the X chromosome and is one of the few X chromosome genes that escapes inactivation (∼15% in women, ∼3% in mice ^4^),^5^ resulting in gene dosage differences between females and males. Mutations in *KDM6A* can cause a rare genetic disorder called Kabuki syndrome which may be associated with CAKUT ^6^ and which is more severe in males.^7^ KDM6A has also been recently linked to the sex-dependent regulation of proximal tubule cell metabolism.^8^ Until now, however, the extent to which KDM6A mediates sex-dependent differences in kidney biology and adaptation to stress has not been known.

KDM6A, also called ubiquitously transcribed tetratricopeptide repeat on chromosome X (UTX), is an ∼154 kDa protein that shares 97% homology between humans and mice.^9^ It possesses an enzymatic Jumonji C (JmjC) domain that catalyzes the removal of the second and third methyl groups from lysine residue 27 on histone protein H3 (H3K27me3),^9^ a mark of gene repression; as well as 6-8 tetratricopeptide repeat domains, which are important for non-enzymatic effects of KDM6A. As such, KDM6A has both H3K27me3 demethylase dependent actions and H3K27me3 demethylase independent actions.^10–15^ Because *Kdm6a* escapes X chromosome inactivation its levels in kidney tubule cells are twice as high in females as in males.^16^ Males, on the other hand, possess one copy of *KDM6A* expressed on the X chromosome, and one copy of its truncated gametolog *UTY* (also called *KDM6C*) expressed on the Y chromosome. This is unusual because the Y chromosome is gene poor. However it is not unique. Humans possess 17 homologous X-Y pairs.^15^ The preservation of a Y chromosome-expressed paralog, with at least some overlap of function,^17, 18^ implies evolutionary pressure to maintain compensation for gene dosage, highlighting that the X chromosome-expressed gene likely performs essential homeostatic functions.

We recently reported that deletion of KDM6A from kidney tubule epithelial cells of male mice is largely inconsequential under normal conditions, and causes subtle, albeit non-negligible changes when the kidney is stressed by ureteric ligation (unilateral ureteral obstruction, UUO).^16^ This led us to question whether KDM6A has a non-essential role in tubule cell homeostasis or whether males are able to compensate for KDM6A absence.^16^ To address this question, in the present study we compared the consequences of tubule cell knockout of KDM6A in female mice and in male mice after UUO. Unexpectedly, we observed that absence of KDM6A from tubule cells of female mice (but not males) induces spontaneous glucosuria under homeostatic conditions, and a Fanconi renotubular syndrome-like picture when the kidney is placed under added stress. We went on to explore the role that KDM6A plays in maintaining tubule epithelial cell homeostasis in females.

## METHODS

### Animals

*Pax8*-Cre^+^ mice that express Cre recombinase controlled by the *Paired box 8* (Pax8) promoter (strain 028196, B6.129P2(Cg)-*Pax8*^tm^^1.1(cre)Mbu^/J) ^19^ were from The Jackson Laboratory (Bar Harbor, ME) and *Kdm6a*^fl/fl^ mice, that possess loxP sites flanking exon 24 of the *Kdm6a* gene,^20^ were provided by Dr. Remy Bosselut (Center for Cancer Research, National Cancer Institute, National Institutes of Health, Bethesda, MD). Using immunohistochemistry and in situ hybridization, we have previously reported that Pax8 is expressed throughout the kidney tubules.^16^ *Pax8*-Cre^+^ mice (KDM6A^Ctrl^) were bred with *Kdm6a*^fl/fl^ mice to generate tubule epithelial cell-specific KDM6A knockout mice (KDM6A^TubKO^; males *Pax8*-Cre^+^*Kdm6a*^fl/Y^, females *Pax8*-Cre^+^*Kdm6a*^fl/fl^). Male and female KDM6A^Ctrl^ and KDM6A^TubKO^ mice underwent sham or UUO surgery at 8-12 weeks of age as previously described.^16^ Briefly, mice were anesthetized with 2% isoflurane, an incision was made in the left flank, and the left ureter was occluded using two 5-0 silk sutures. Sham mice underwent the same procedure without ligation of the left ureter. Slow-release buprenorphine (0.5 mg/kg subcutaneously) was administered pre-operatively for analgesia. The mice were followed for 14 days after surgery. At the end of the study period, mice were individually housed in metabolic cages for 24 hours to measure water consumption and urine output. In sub-groups of mice, fasting blood glucose was measured by OneTouch UltraMini (LifeScan Canada, Burnaby, British Columbia, Canada), and HbA_1c_ was measured using A1cNow (PTS Diagnostics, Whitestown, IN). Urine electrolytes and metabolites were measured at The Centre for Phenogenomics (Toronto, Ontario, Canada). After harvesting, mouse kidneys were immersed in 10% neutral buffered formalin, routinely processed, and embedded in paraffin; flash frozen in liquid nitrogen and stored at -80°C; or fixed in 2.5% glutaraldehyde for later assessment by transmission electron microscopy. All experimental procedures adhered to the guidelines of the Canadian Council on Animal Care and were approved by the St. Michael’s Hospital Animal Care Committee (ACC 181).

### Primary tubule epithelial cells

Tubule epithelial cells were isolated from wildtype and KDM6A^TubKO^ mouse kidneys following the protocol by Ding et al,^21^ and as we have previously described.^16, 22^ We have previously characterized these cells for their expression of CD10 and aquaporin 1.^22^

### Immunoblotting

Immunoblotting was performed on homogenized kidney tissues and cell lysates using the following antibodies, each at 1:1000 dilution: KDM6A (catalog no. 33510, clone D3Q1I, lot no. 2, Cell Signaling Technology, Danvers, MA), SGLT2 (catalog no. 24654-1-AP, lot no. 00161680, Thermo Fisher Scientific, Waltham, MA), GLUT2 (catalog no. 20436-1-AP, lot no. 00162787, Thermo Fisher Scientific), H3K27me3 (catalog no. 9733s, lot no. 14, Cell Signaling Technology), histone H3 (catalog no. 14269s, lot no. 2, Cell Signaling Technology) and GAPDH (catalog no. 2118, clone 14C10, lot no. 14, Cell Signaling Technology). Densitometry was performed using ImageJ software version 1.53 (NIH, Bethesda, MD; https://imagej.net/ij).

### Histology

For quantitation of interstitial fibrosis, mouse kidney sections were stained with picrosirius red (ab246832; Abcam, Waltham, MA) and digitized (Axio Scan.z1, Carl Zeiss Canada Ltd., Toronto, ON, Canada), and the proportional area positively staining red was analyzed using HALO (Indica Labs, Albuquerque, NM). For visualization of tubule cell integrity, kidney sections were stained with H&E and for additional visualization of interstitial fibrosis, kidney sections were stained with Masson’s trichrome by members of the Pathology Research Program at Toronto General Hospital (Toronto, Ontario, Canada). Glomerular architecture was assessed following staining of kidney sections with Periodic acid-Schiff.

### Quantitative real-time RT-PCR

RNA was isolated from mouse kidney tissue using TRIzol Reagent (Thermo Fisher Scientific), and cDNA was reverse transcribed from 2 μg RNA using Applied Biosystems™ High-Capacity cDNA Reverse Transcription Kit (catalog no. 4368814; lot no. 2890929, Thermo Fisher Scientific). cDNA was diluted 1:20 before use. Primers were designed and validated using Primer-BLAST (http://www.ncbi.nlm.nih.gov/tools/primer-blast/) or Harvard Primer Bank (https://pga.mgh.harvard.edu/primerbank/), and they were synthesized by Integrated DNA Technologies (Coralville, IA). Primer sequences are listed in S. I. Table 1. SYBR Green-based qRT-PCR was performed on a QuantStudio 7 Flex Real-Time PCR System (Thermo Fisher Scientific) and data were analyzed using the comparative CT (ΔΔCT) method.

**Table 1.** Urine electrolytes and metabolites in male KDM6A^Ctrl^ and KDM6A^TubKO^ mice 14 days after sham surgery or unilateral ureteral obstruction (UUO).

|  | KDM6A<sup>Ctrl</sup> sham | KDM6A<sup>Ctrl</sup> UUO | KDM6A<sup>TubKO</sup> sham | KDM6A<sup>TubKO</sup> UUO |
| --- | --- | --- | --- | --- |
| *n* | 10 | 10 | 12 | 13 |
| BUN (mg/24h) | 14.9 ± 6.6 | 16.9 ± 9.4 | 13.8 ± 9.0 | 19.4 ± 7.7 |
| Total protein<br>(mg/24h) | 3.0 ± 1.3 | 4.5 ± 3.7 | 2.3 ± 1.7 | 3.0 ± 1.4 |
| Creatinine<br>(mg/24h) | 0.25 ± 0.14 | 0.24 ± 0.14 | 0.21 ± 0.11 | 0.27 ± 0.08 |
| Albumin<br>(mg/24h) | 0.002 ± 0.005 | 0.007 ± 0.008 | 0.005 ± 0.006 | 0.006 ± 0.008 |
| Sodium<br>(mmol/24h) | 0.09 ± 0.04 | 0.07 ± 0.05 | 0.06 ± 0.04 | 0.09 ± 0.03 |
| Potassium<br>(mmol/24h) | 0.14 ± 0.06 | 0.14 ± 0.07 | 0.10 ± 0.07 | 0.16 ± 0.06 |
| Chloride<br>(mmol/24h) | 0.10 ± 0.05 | 0.08 ± 0.05 | 0.07 ± 0.05 | 0.11 ± 0.05 |
| Phosphate<br>(mg/24h) | 1.3 ± 0.9 | 1.3 ± 1.0 | 1.0 ± 0.7 | 1.5 ± 0.7 |
| Calcium<br>(mg/24h) | 0.06 ± 0.02 | 0.08 ± 0.05 | 0.06 ± 0.03 | 0.10 ± 0.03<sup>ab</sup> |
| Glucose<br>(mg/24h) | 0.21 ± 0.10 | 0.19 ± 0.09 | 0.15 ± 0.14 | 0.22 ± 0.10 |

Values are mean ± S.D.. <sup>a</sup>*P* < 0.05 vs. KDM6A<sup>Ctrl</sup> sham, <sup>b</sup>*P* < 0.05 vs. KDM6A<sup>TubKO</sup> sham by one-way ANOVA followed by Tukey's post-hoc test.

### RNAscope in situ hybridization

RNAscope in situ hybridization (Advanced Cell Diagnostics, Hayward, CA) was performed on kidney sections from male mice according to the manufacturer’s instructions,^23, 24^ and using the following probesets: *Kdm6a* (catalog no. 456961, lot no. 24163A), *Uty* (catalog no. 451741, lot no. 24271D), *dapB* (catalog no. 310043, lot no. 2017575). Tubule epithelial *Kdm6a* and *Uty* mRNA was quantified by counting RNAscope puncta within kidney tubule cells in six randomly selected kidney cortical fields (x 400 magnification) per mouse, in a masked manner on a Nikon Eclipse e200 microscope (Nikon Instruments Inc., Melville, NY).

### Spatial transcriptomics

Spatial transcriptomics was performed at the Princess Margaret Genomics Centre (PMGC; Toronto, Ontario, Canada) using the 10x Genomics Visium HD Spatial Gene Expression system. Briefly, formalin-fixed paraffin-embedded (FFPE) mouse kidney tissues were sectioned at 5 µm thickness from four female KDM6A^Ctrl^ mice and four female KDM6A^TubKO^ mice. Sections were mounted onto a Fisherbrand SuperFrost Plus slide (PN# 12-550-15, Fisher Scientific, Mississauga, Ontario, Canada) with the four KDM6A^Ctrl^ sections in one area and the four KDM6A^TubKO^ sections in another area (each area being approximately equivalent to the 6.5 mm x 6.5 mm capture area on a Visium slide), according to the manufacturer’s instructions (https://cdn.10xgenomics.com/image/upload/v1756507543/support-documents/CG000684_VisiumHDFFPETissuePrepHandbook_RevD.pdf). Tissue sections were baked, deparaffinized using xylene and graded ethanol washes, and stained with H&E for histological visualization and imaging. Spatial transcriptomics libraries were prepared according to the 10x Genomics Visium HD Spatial Gene Expression Reagent Kits instructions (https://cdn.10xgenomics.com/image/upload/v1756507345/support-documents/CG000685_VisiumHD_GeneExpression_UserGuide_RevD.pdf). Briefly, following coverslip removal, slides were assembled in the Visium cassette and destained and decrosslinked, prior to probe hybridization using the mouse whole transcriptome probe panel, and probe ligation. The Visium HD slide was thawed, washed and equilibrated before placement in the cassette with equilibration mix. The tissue slide was removed from the cassette, stained with eosin, washed and loaded with the Visium slide into the Visium CytAssist instrument for probe release and capture. Single stranded ligation products were released from the tissue upon RNase treatment and captured on the Visium slide before removal from the Visium CytAssist. Spatially barcoded ligation products were generated by extending ligation products with the addition of Spatial Barcode, unique molecular identifier (UMI), and partial Read 1 primer. Spatially barcoded ligation products were then carried forward for library preparation. qPCR was performed on the pre-Amplification products to determine the number of PCR cycles for library preparation (Cq: KDM6A^Ctrl^ 8.44; KDM6A^TubKO^ 8.63), and pre-Amplification products were amplified at 10 cycles. Libraries were sequenced using a NovaSeq X (paired-end, dual indexed sequencing, 500,000,000 reads; Read1T 43 cycles, i7 index 10 cycles, i5 index 10 cycles, Read 2S 50 cycles).

Visium HD libraries were processed using the Space Ranger count pipeline (10x Genomics). Sequencing reads were aligned to the mm10 mouse reference genome, and transcript counts were quantified using the probe-based gene expression approach. Cell segmentation was performed by Space Ranger using a custom implementation of StarDist,^25^ a deep learning model for instance segmentation of cell nuclei applied to the H&E tissue image. Per-cell UMI counts were used for all downstream analyses. Tissue sections within each capture area were computationally separated using density-based spatial clustering (DBSCAN) applied to the centroid coordinates of segmented cells, with parameters eps = 400 and minPts = 50. This yielded eight independent spatial objects (WT_1–WT_4, KO_1–KO_4), each annotated with a group label (WT for KDM6A^Ctrl^ or KO for KDM6A^TubKO^) and a unique sample_id used for all pseudobulk analyses. All eight section objects were merged into a single Seurat object ^26^ for quality control and downstream analyses. The percentage of mitochondrial reads was computed per cell. High quality cells were retained based on the following criteria: (1) number of detected genes > 150 and < 2,200; (2) total UMI count > 400 and < 5,000; (3) percentage of mitochondrial reads < 25%. Filtered cells were normalized per sample using SCTransform, regressing out the percentage of mitochondrial reads. Samples were then integrated using the RPCAIntegration procedure in Seurat. Uniform Manifold Approximation and Projection (UMAP) dimensionality reduction and nearest-neighbor graph construction were performed based on the first 30 principal components from the RPCA embedding, followed by graph-based clustering using the FindClusters function at resolution = 0.5. Cluster-specific marker genes were identified using FindMarkers with the Wilcoxon rank-sum test (cutoffs: log_₂_ fold-change > 0.5, Benjamini-Hochberg adjusted *P* value < 0.05, upregulated genes only). Clusters were annotated to kidney cell-types by manual curation based on canonical marker gene expression. Per cell-type differential expression testing between KO and WT conditions was conducted using a pseudobulk approach. Raw counts were summed across all cells sharing the same sample and cell-type annotation using the AggregateExpression function in Seurat. Pseudobulk columns with fewer than 50 cells contributing were excluded from analysis. Lowly expressed genes with an average read count lower than 10 across all retained samples within a given cell-type were also excluded. The cell-type LOH (IOM) was excluded from differential expression testing due to insufficient representation across samples. Raw pseudobulk counts were normalized using edgeR’s TMM with singleton pairing algorithm ^27^ and were then transformed to log_2_-counts per million (log_2_CPM) using the voomLmFit function implemented in the R package limma.^28^ A cell-type specific linear model was fitted for each KO versus WT contrast using makeContrasts and eBayes with robust estimation. Nominal *P* values were corrected for multiple testing using the Benjamini-Hochberg method. Differentially expressed genes were defined as those with an adjusted *P* value < 0.05 and log_₂_ fold-change > 0.5. Gene set enrichment analysis (GSEA) based on a pre-ranked gene list (ranked by moderated t-statistic) was performed using the R package fgsea ^29^ against the MSigDB Gene Ontology Biological Process (GO:BP) gene sets for *Mus musculus*. Pathways with a minimum gene set size of 5 and a Benjamini-Hochberg adjusted *P* value < 0.05 were considered significant.

Data are deposited to Gene Expression Omnibus (accession number GSE336198).

### Transmission electron microscopy

Transmission electron microscopy was performed using a Hitachi HT7800 (Hitachi High-Tech Europe, Abingdon, United Kingdom) at Electron Microscopy Research Services (Newcastle University, Newcastle, UK) on kidney cortical tissue from four female KDM6A^Ctrl^ and four female KDM6A^TubKO^ mice. Mitochondrial roundness, aspect ratio and circularity were measured in five proximal tubule cross sections from each mouse using ImageJ according to the method described by Faitg et al.^30^

### Untargeted metabolomics

Untargeted metabolomics was performed on kidney tissue from four female KDM6A^Ctrl^ mice and four female KDM6A^TubKO^ mice at the Segal Cancer Proteomics Centre, Lady Davis Institute for Medical Research, Jewish General Hospital and McGill University (Montreal, QC, Canada). Briefly, mouse frozen kidney samples were reconstituted in 200 µL of 80% MeCN containing 1 µmol/L caffeine-d9 as an internal standard. The samples were homogenized, sonicated for 30 min and centrifuged at 21000 x g for 1 min at room temperature (23°C). One hundred and eighty microlitres of supernatants were lyophilized, reconstituted in 180 µl 0.1% FA, sonicated for one minute and centrifuged at 21000 x g for 1 min at room temperature. One hundred microlitres of each sample supernatant were used for LC-MS/MS analysis. Pooled samples from each group were made by combining 10 µl of each sample of the same group (Total). LC-MS analyses were performed using Vanquish UPLC with 0-100% 20 min gradient (0.1% FA - 0.1%FA/MeCN) at 200 µL/min flow rate on an Agilent InfinityLab Poroshell 120 Aq-C18 column 2.1 x 150 mm, 2.7 µm (RP-LC-MS). Ten microlitre injection volumes were used for each run. Mass spectra were acquired using a Thermo Orbitrap QE+ spectrometer in positive and negative modes in separate runs. Data for each LC/acquisition type were analyzed using Compound Discoverer 3.3 (Thermo). KEGG pathway analysis was performed using the Functional Analysis tool of MetaboAnalyst 6.0 ^31^ with input data: p_value, m/z, fold change and retention time with *P* < 0.05 as cut off, along with the lipid class and lipid subclass filters for lipidomics. This study is available at the NIH Common Fund’s National Metabolomics Data Repository (NMDR) website, the Metabolomics Workbench,^32^ https://www.metabolomicsworkbench.org where it has been assigned Study ID ST005038. The data can be accessed directly via the Project DOI: http://dx.doi.org/10.21228/M8D86N.

### Statistics

Data are expressed as means ± S.D.. Animals were randomly allocated to sham or UUO groups. Analyses of data were performed in a masked manner where feasible. Statistical analyses were performed using GraphPad Prism 10 for macOS (GraphPad Software Inc., San Diego, CA). Comparison between two groups was by Student *t* test and between multiple groups was by one-way ANOVA followed by Tukey post-test unless otherwise stated. For multiple comparisons, post hoc testing was only conducted if F in the analysis of variance achieved *P <* 0.05. All analyses were two tailed. *P <* 0.05 was considered statistically significant.

## RESULTS

### Knockout of KDM6A from tubule cells augments kidney fibrosis in female mice with UUO

After we had confirmed absence of KDM6A from primary cultured tubule epithelial cells (TECs) isolated from male and female KDM6A^TubKO^ mouse kidneys (Fig. 1A), KDM6A^Ctrl^ and KDM6A^TubKO^ mice were randomized to undergo sham or UUO surgery and they were followed for 14 days. Interstitial fibrosis, determined by picrosirius red staining, was increased in male mice 14 days after UUO and it was unaffected by tubule cell-specific KDM6A knockout (Fig. 1B). In contrast, tubule cell-specific knockout of KDM6A in female mice augmented interstitial fibrosis 14 days after UUO (Fig. 1C).

**Figure 1.**
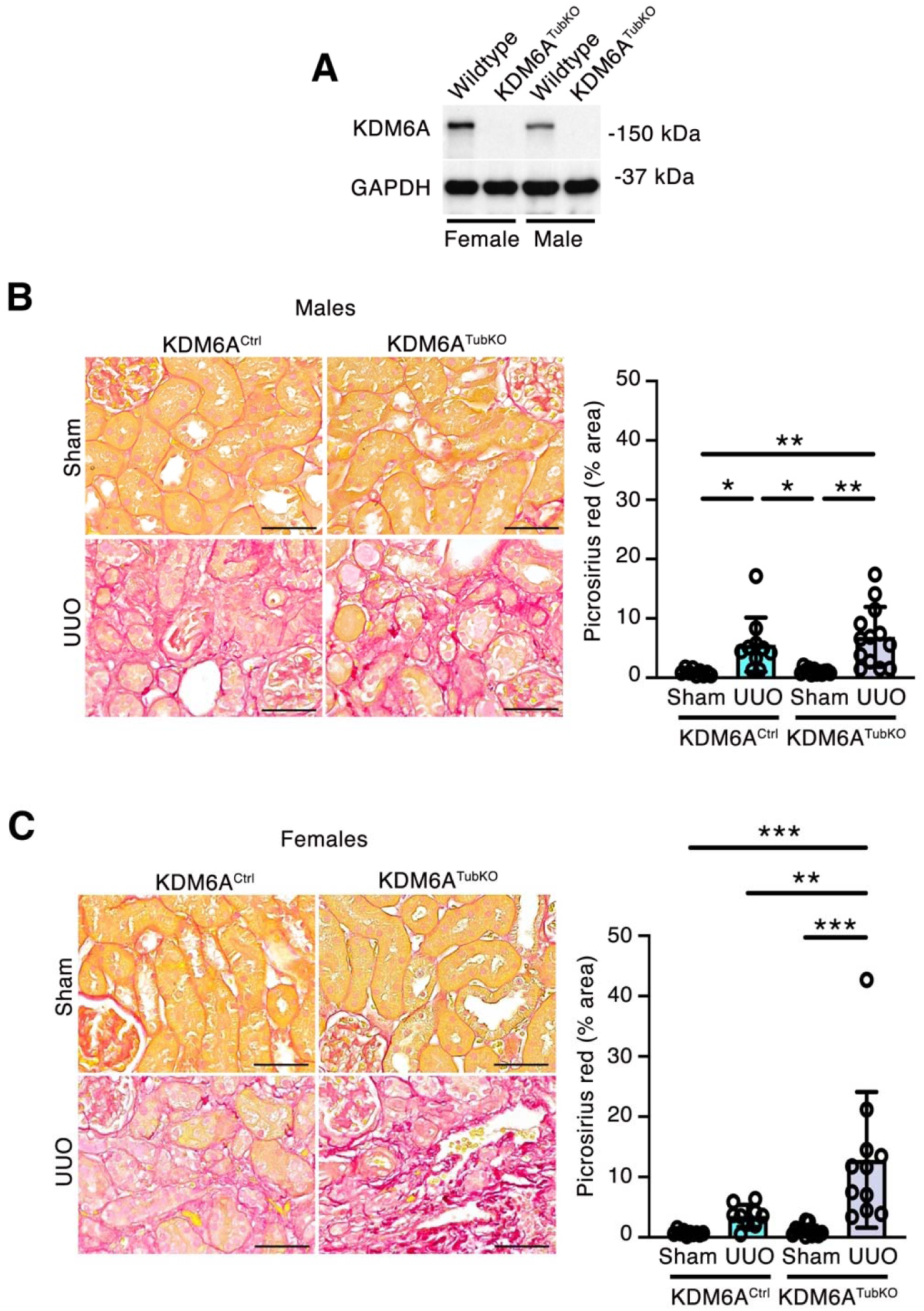
Kidney fibrosis is augmented in female KDM6A^TubKO^ mice but not in males. (A) Immunoblotting for KDM6A in male and female wildtype and KDM6A^TubKO^ tubule epithelial cells showing absence of KDM6A in male and female KDM6A^TubKO^ mice. Representative of n=3/group. (B and C) Male and female KDM6A^Ctrl^ and KDM6A^TubKO^ mice underwent sham or unilateral ureteral obstruction (UUO) surgery and were followed for 14 days. Representative picrosirius red staining and quantitation of staining in male (B) and female (C) mice. Original magnification x 400. Scale bar = 50 µm. (B) Male KDM6A^Ctrl^ sham n=9, male KDM6A^Ctrl^ UUO n=10, male KDM6A^TubKO^ sham n=12, male KDM6A^TubKO^ UUO n=13. (C) Female KDM6A^Ctrl^ sham n=10, female KDM6A^Ctrl^ UUO n=9, female KDM6A^TubKO^ sham n=12, female KDM6A^TubKO^ UUO n=11. Values are mean ± S.D.. \**P* < 0.05, \*\**P* < 0.01, \*\*\**P* < 0.001 by one-way ANOVA followed by Tukey’s post-test.

### Knockout of KDM6A from tubule cells increases water consumption and urine output in female mice with UUO

Figure 2 shows the body weights (Figs. 2A and B), left (obstructed) kidney weights (Figs. 2C-F), right (unobstructed) kidney weights (Figs. 2G and H), water consumption (Figs. 2I and J) and urine output (Figs. 2K and L) in male and female KDM6A^Ctrl^ and KDM6A^TubKO^ mice 14 days after sham or UUO surgery. Body weights were slightly lower in KDM6A^TubKO^ mice than in KDM6A^Ctrl^ mice. As expected, the weights of the left (obstructed) kidneys were increased in male and female UUO mice (Figs. 2C-F), and they were unaffected by tubule cell-specific KDM6A knockout in either males or females (Figs. 2C-F). There was also compensatory hypertrophy of the right (unobstructed) kidneys in both male and female UUO mice that was also unaffected by KDM6A knockout (Figs. 2G and H). Water consumption and urine output were unaltered by either UUO or tubule cell-specific KDM6A knockout in male mice (Figs. 2I and K). Unexpectedly, however, both water intake and urine output were numerically higher in female sham-operated KDM6A^TubKO^ mice than KDM6A^Ctrl^ mice, and they were significantly increased in female KDM6A^TubKO^ mice following UUO (Figs. 2J and L).

**Figure 2.**
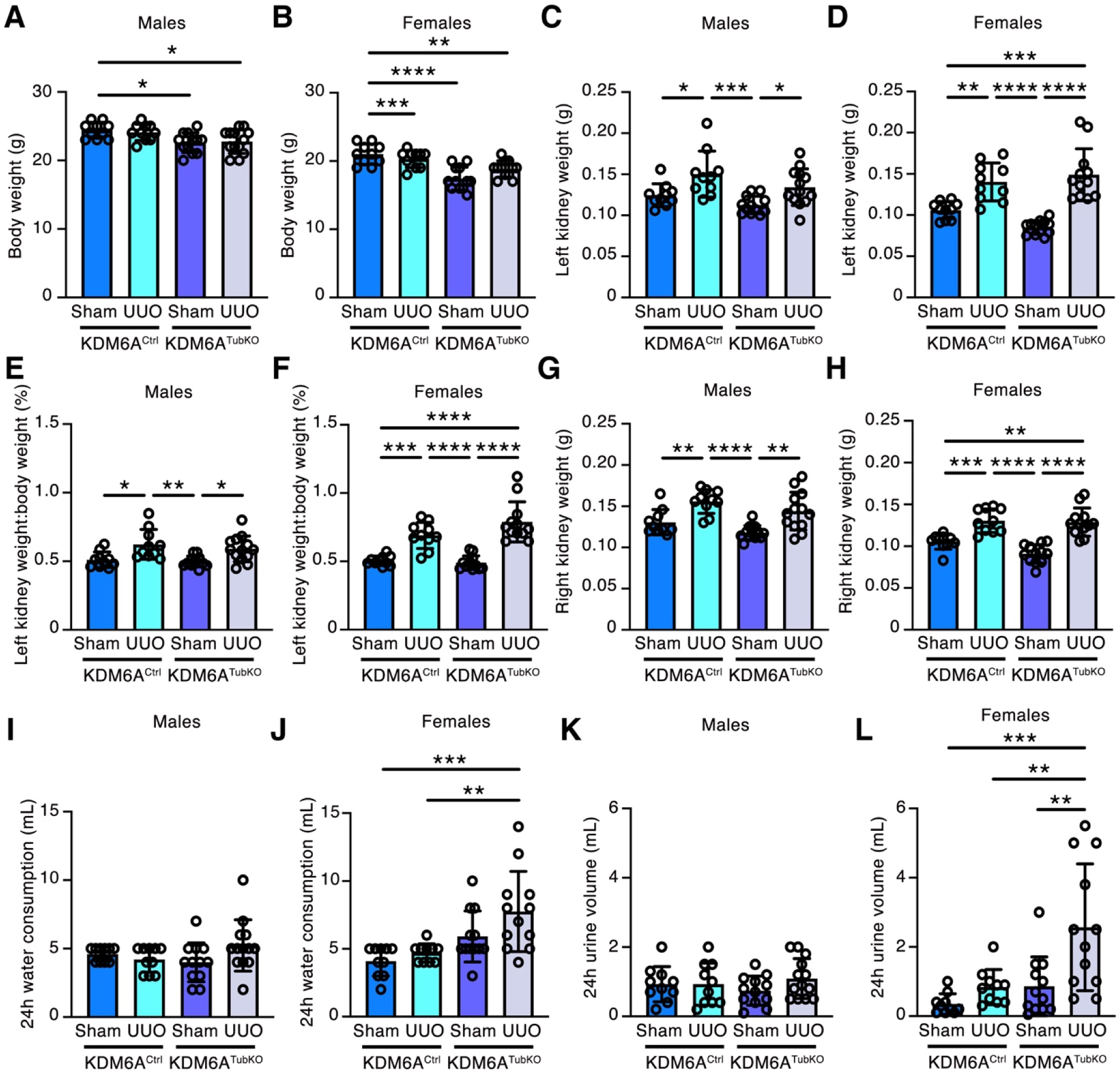
Water consumption and urine volume are increased in female but not male KDM6A^TubKO^ mice with unilateral ureteral obstruction (UUO). Male and female KDM6A^Ctrl^ and KDM6A^TubKO^ mice underwent sham or UUO surgery and were followed for 14 days. (A and B) Body weight in male (A) and female (B) mice. (C and D) Left (obstructed) kidney weight in male (C) and female (D) mice. (E and F) Left kidney weight:body weight ratio in male (E) and female (F) mice. (G and H) Right kidney weight in male (G) and female (H) mice. (I and J) 24h water consumption in male (I) and female (J) mice. (K and L) 24h urine volume in male (K) and female (L) mice. Male KDM6A^Ctrl^ sham n=10, male KDM6A^Ctrl^ UUO n=10, male KDM6A^TubKO^ sham n=12, male KDM6A^TubKO^ UUO n=13; female KDM6A^Ctrl^ sham n=10, female KDM6A^Ctrl^ UUO n=10, female KDM6A^TubKO^ sham n=12, female KDM6A^TubKO^ UUO n=12. Values are mean ± S.D.. \**P* < 0.05, \*\**P* < 0.01, \*\*\**P* < 0.001, \*\*\*\**P* < 0.0001 by one-way ANOVA followed by Tukey’s post-test.

### Knockout of KDM6A from tubule cells causes glucosuria in female but not male mice

Intrigued by the increase in urine volume in female KDM6A^TubKO^ mice, especially after UUO, we next measured urine electrolytes and metabolites (Tables 1 and 2). Most notably, we observed that either sham or UUO female KDM6A^TubKO^ mice developed overt glucosuria (Table 2), whereas male KDM6A^TubKO^ mice did not (Table 1). Twenty four hour urine excretion of BUN, creatinine, albumin, sodium, potassium, phosphate and calcium were all also increased in female KDM6A^TubKO^ mice under UUO conditions (Table 2). Thus, glucosuria and a Fanconi renotubular syndrome-like picture in female KDM6A^TubKO^ mice after UUO likely underlies the polyuria these mice develop. Because the ureter exiting the left kidney is ligated with UUO, this increase in solute excretion arose from the unobstructed (right) kidney. To exclude diabetes mellitus as a cause of the glucosuria, in separate cohorts, we measured fasting blood glucose levels and HbA_1c_ levels in male and female KDM6A^Ctrl^ and KDM6A^TubKO^ mice, observing no difference in glycemia despite the development of glucosuria in female KDM6A^TubKO^ mice (Figs. 3A-D). Genetic mutations of *SLC5A2*, the gene that encodes SGLT2, are the most common cause of heritable renal glucosuria.^33^ Accordingly, we next performed qRT-PCR and immunoblotting for *Slc5a2*/SGLT2 in the kidneys of sham and UUO male and female KDM6A^Ctrl^ and KDM6A^TubKO^ mice (studying the unobstructed kidneys). Whereas *Slc5a2* mRNA levels were unaltered in male mice (Fig. 3E), they were numerically lower in female KDM6B^TubKO^ mice and they were significantly diminished in the unobstructed kidneys of female KDM6B^TubKO^ mice after UUO (Fig. 3F). SGLT2 protein levels were similarly unaltered in males (Fig. 3G), whereas they were diminished in female KDM6B^TubKO^ mice, even under sham conditions (Fig. 3H). In contrast, protein levels of the basolateral glucose transporter, GLUT2 were unaltered across groups (S. I. Fig. 1A). H3K27me3 levels were unaltered in either male or female KDM6A^TubKO^ mice under either sham or UUO conditions (S. I. Fig. 1B).

**Figure 3.**
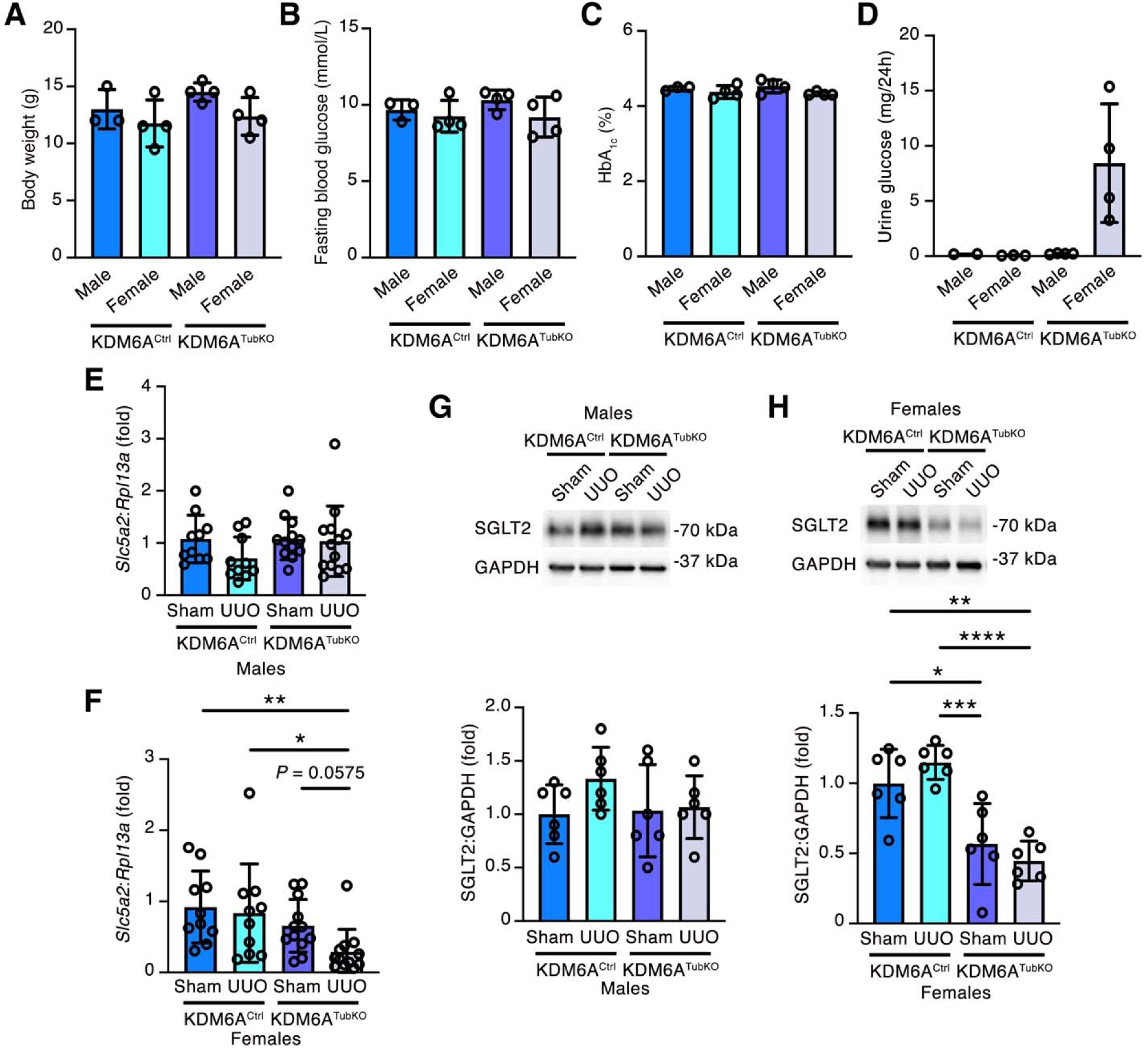
SGLT2 protein levels are diminished in the kidneys of female KDM6A^TubKO^ mice. (A-D) Body weight (A), fasting blood glucose (B), HbA_1c_ (C) and 24h urine glucose excretion (D) in male and female KDM6A^Ctrl^ and KDM6A^TubKO^ mice. Male KDM6A^Ctrl^ (n=3, except (D) n=2), female KDM6A^Ctrl^ (n=4, except (D) n=3), male KDM6A^TubKO^ (n=4), female KDM6A^TubKO^ (n=4). (E and F) qRT-PCR for *Slc5a2* in male (E) and female (F) mice. Male and female KDM6A^Ctrl^ and KDM6A^TubKO^ mice underwent sham or UUO surgery and were followed for 14 days. (E) Male KDM6A^Ctrl^ sham n=10, male KDM6A^Ctrl^ UUO n=10, male KDM6A^TubKO^ sham n=12, male KDM6A^TubKO^ UUO n=13. (F) Female KDM6A^Ctrl^ sham n=10, female KDM6A^Ctrl^ UUO n=10, female KDM6A^TubKO^ sham n=12, female KDM6A^TubKO^ UUO n=12. (G and H) Immunoblotting for SGLT2 in male (G) and female (H) mice. n=6/group. Values are mean ± S.D.. \**P* < 0.05, \*\**P* < 0.01, \*\*\**P* < 0.001, ****P < 0.0001 by one-way ANOVA followed by Tukey’s post-test.

**Table 2.** Urine electrolytes and metabolites in female KDM6A^Ctrl^ and KDM6A^TubKO^ mice 14 days after sham surgery or unilateral ureteral obstruction (UUO).

|  | KDM6A<sup>Ctrl</sup> sham | KDM6A<sup>Ctrl</sup> UUO | KDM6A<sup>TubKO</sup> sham | KDM6A<sup>TubKO</sup> UUO |
| --- | --- | --- | --- | --- |
| *n* | 8 | 10 | 11 | 12 |
| BUN (mg/24h) | 8.4 ± 6.7 | 16.6 ± 10.1 | 17.7 ± 15.4 | 27.1 ± 15.9<sup>a</sup> |
| Total protein (mg/24h) | 0.5 ± 0.3 | 0.7 ± 0.3 | 0.9 ± 0.8 | 1.4 ± 1.4 |
| Creatinine (mg/24h) | 0.12 ± 0.09 | 0.15 ± 0.06 | 0.14 ± 0.09 | 0.26 ± 0.09<sup>ab</sup> |
| Albumin (mg/24h) | 0.003 ± 0.003 | 0.008 ± 0.006 | 0.019 ± 0.019 | 0.052 ± 0.033<sup>cd</sup> |
| Sodium (mmol/24h) | 0.04 ± 0.02 | 0.06 ± 0.04 | 0.06 ± 0.04 | 0.10 ± 0.05<sup>a</sup> |
| Potassium (mmol/24h) | 0.06 ± 0.05 | 0.12 ± 0.07 | 0.14 ± 0.13 | 0.21 ± 0.12<sup>e</sup> |
| Chloride (mmol/24h) | 0.04 ± 0.03 | 0.08 ± 0.05 | 0.10 ± 0.09 | 0.13 ± 0.09 |
| Phosphate (mg/24h) | 0.5 ± 0.4 | 1.2 ± 0.8 | 0.9 ± 1.1 | 2.2 ± 1.0<sup>be</sup> |
| Calcium (mg/24h) | 0.05 ± 0.03 | 0.10 ± 0.05<sup>a</sup> | 0.08 ± 0.05 | 0.12 ± 0.05<sup>e</sup> |
| Glucose (mg/24h) | 0.18 ± 0.15 | 0.49 ± 0.56 | 18.71 ± 19.24<sup>df</sup> | 25.71 ± 22.23<sup>fg</sup> |

Values are mean ± S.D.. <sup>a</sup>*P* < 0.05 vs. KDM6A<sup>Ctrl</sup> sham, <sup>b</sup>*P* < 0.05 vs. KDM6A<sup>TubKO</sup> sham, <sup>c</sup>*P* < 0.0001 vs. KDM6A<sup>Ctrl</sup> sham, <sup>d</sup>*P* < 0.05 vs. KDM6A<sup>Ctrl</sup> UUO, <sup>e</sup>*P* < 0.01 vs. KDM6A<sup>Ctrl</sup> sham, <sup>f</sup>*P* < 0.001 vs. KDM6A<sup>Ctrl</sup> sham, <sup>g</sup>*P* < 0.01 vs. KDM6A<sup>Ctrl</sup> UUO by Kruskal-Wallis test followed by Dunn's post-test.

### Knockout of KDM6A from tubule cells in female mice causes tubular epithelial cell vacuolization and atrophy, and focal interstitial lymphoid infiltration and fibrosis

Next, we surveyed kidney structure histologically, studying the unobstructed kidneys in the UUO mice as well as the normal kidneys of sham-operated mice. H&E staining revealed that female KDM6A^TubKO^ mice developed focal areas of tubule epithelial vacuolization, tubule atrophy and lymphoid cell infiltration both under sham and UUO conditions, whereas tubule cell architecture appeared normal in male KDM6A^TubKO^ mice (Figs. 4A and B). These interstitial changes in female KDM6A^TubKO^ mice were accompanied by areas of collagen deposition, most notable in the unobstructed kidneys of female KDM6A^TubKO^ mice after UUO, which was again not observed in male KDM6A^TubKO^ mice (Figs. 4C and D). Glomerular architecture was preserved in both male and female KDM6A^TubKO^ mice (S. I. Fig. 2). Paralleling the changes observed in focal interstitial fibrosis, mRNA levels of the proinflammatory and profibrotic cytokine *Tnf*, whilst marginally elevated in unobstructed kidneys of male KDM6A^TubKO^ mice after UUO, were markedly increased in female KDM6A^TubKO^ mice after UUO (Figs. 4E and F). Thus, absence of KDM6A from tubule epithelial cells in female (but not male) mice causes tubule cell injury that manifests as focal vacuolization, reduced SGLT2 expression, glucosuria and decreased ability to withstand added stress caused by obstruction of urinary outflow from the contralateral kidney.

**Figure 4.**
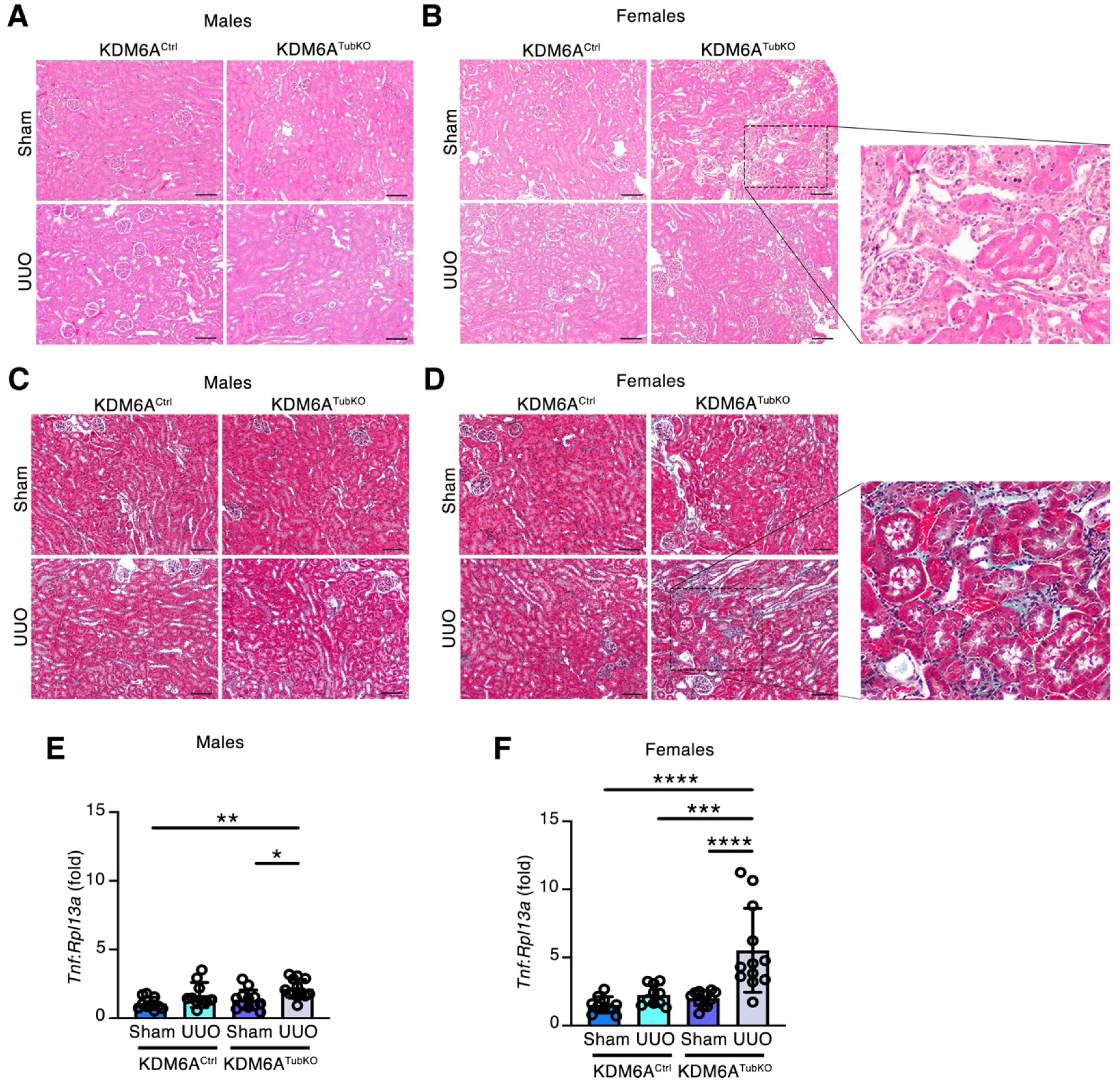
Knockout of KDM6A from tubule cells causes tubule cell vacuolization in female mice, and interstitial fibrosis and *Tnf* upregulation in unobstructed kidneys of female mice after UUO. (A and B) Representative H&E staining of kidneys of male (A) and female (B) KDM6A^Ctrl^ and KDM6A^TubKO^ mice 14 days after sham surgery or of the unobstructed (right) kidneys of mice 14 days after unilateral ureteral obstruction (UUO). n=4/group. Original magnification x 100. Scale bar = 100 µm. (C and D) Masson’s trichrome staining of kidneys of male (C) and female (D) KDM6A^Ctrl^ and KDM6A^TubKO^ mice 14 days after sham surgery or of the unobstructed (right) kidneys of mice 14 days after UUO. n=4/group. Original magnification x 100. Scale bar = 100 µm. In B and D the zoomed in images are enlargements of the dashed areas. (E and F) qRT-PCR for *Tnf* in the kidneys of male (E) and female (F) KDM6A^TubKO^ mice 14 days after sham surgery or of the unobstructed (right) kidneys of male and female KDM6A^TubKO^ mice 14 days after UUO. (E) Male KDM6A^Ctrl^ sham n=10, male KDM6A^Ctrl^ UUO n=10, male KDM6A^TubKO^ sham n=12, male KDM6A^TubKO^ UUO n=13. (F) Female KDM6A^Ctrl^ sham n=10, female KDM6A^Ctrl^ UUO n=10, female KDM6A^TubKO^ sham n=12, female KDM6A^TubKO^ UUO n=12. Values are mean ± S.D.. \**P* < 0.05, \*\**P* < 0.01, \*\*\**P* < 0.001, ****P < 0.0001 by one-way ANOVA followed by Tukey’s post-test.

### *Uty* and *Kdm6a* are expressed at equivalent levels in tubule cells of male mice and are each upregulated with UUO

Given that glucosuria, tubule injury and stress intolerance were apparent in female mice with tubule cell-specific KDM6A deletion, but not in males, we probed for expression of the Y chromosome expressed paralog, *Uty* in the kidneys of male mice. Using RNAscope in situ hybridization, we observed a similar pattern of expression of *Uty* as for *Kdm6a*, transcripts for each being present in multiple different kidney cell-types including tubule epithelial cells (Figs. 5A and B), and both *Kdm6a* and *Uty* being upregulated equivalently in UUO kidneys (Figs. 5A and B).

**Figure 5.**
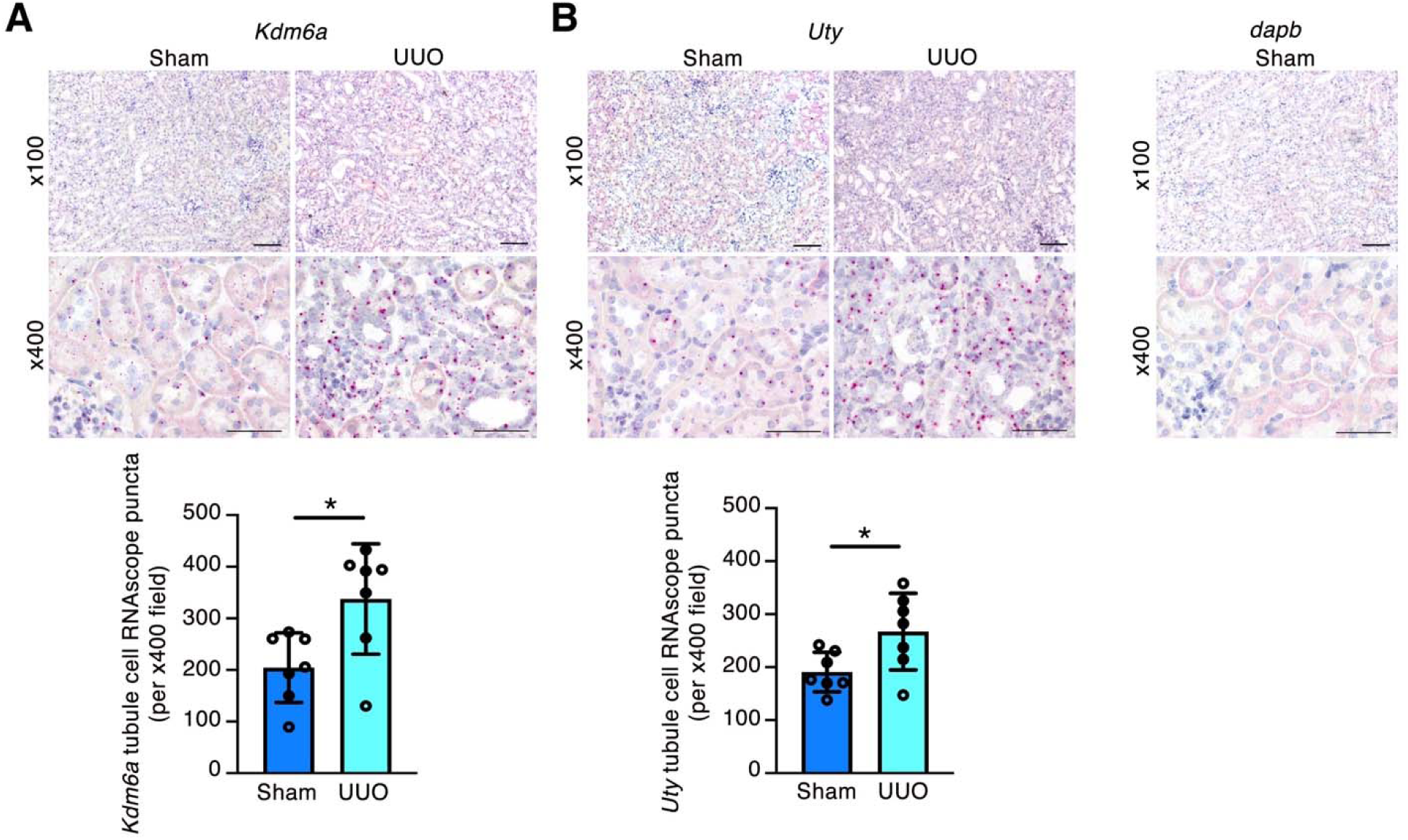
*Kdm6a* and *Uty* exhibit similar patterns of expression in male kidneys, with both transcripts being present in tubule epithelial cells and other cells and both transcripts being similarly upregulated after unilateral ureteral obstruction (UUO). RNAscope in situ hybridization for *Kdm6a* (A) and *Uty* (B) and quantitation of RNAscope puncta in the kidneys of KDM6A^Ctrl^ mice 14 days after sham or UUO (n=7/group). x 100 magnification, scale bar = 100 µm; x 400 magnification, scale bar = 50 µm. The bacterial transcript *dapB* is the negative control. Values are mean ± S.D.. \**P* < 0.05 by two-tailed Student *t* test.

### Knockout of KDM6A induces transcriptional reprogramming in proximal tubule cells of female mice

To determine how knockout of KDM6A affects the transcriptional machinery of tubule epithelial cells, provoking injury and stress intolerance, we next performed spatial transcriptomics of the kidneys of female KDM6A^Ctrl^ and KDM6A^TubKO^ mice (Figs. 6A and B). This analysis encompassed 152,408 cells in KDM6A^Ctrl^ mice and 152,823 cells in KDM6A^TubKO^ mice, detecting 17,858 genes in KDM6A^Ctrl^ mice and 17,690 genes in KDM6A^TubKO^ mice, from four mice for each genotype (S. I. Table 2). Using canonical marker gene expression (S. I. Fig. 3),^34^ we were able to distinguish 16 cell-types: ascending loop of Henle in cortex (ALOH), collecting duct (CD), connecting tubules and collecting ducts (CNT_CD), distal tubules and connecting tubules (DT_CNT), glomerular cells, immune cells, injured proximal tubule (InjPT), injured proximal tubule segment 3 cells (InjPT3), juxtaglomerular cells, loop of Henle in outer medulla (LOH (IOM)), proximal tubule S1 (PTS1) segment, proximal tubule S1S2 (PTS1S2) segment, proximal tubule S2 (PTS2) segment, proximal tubule S3 (PTS3) segment, vasa recta, and vasculature (Figs. 6C and D).

**Figure 6.**
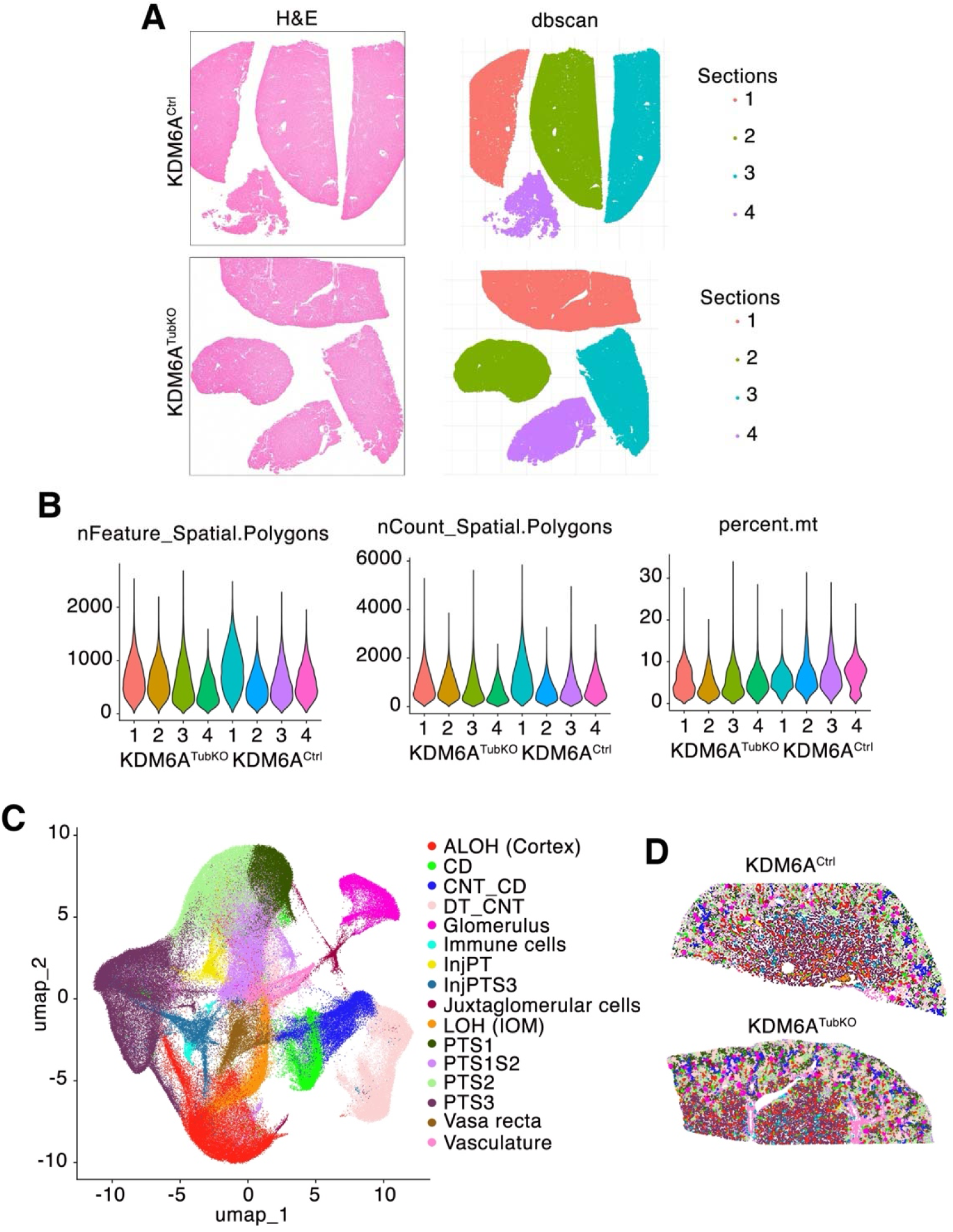
Detection of 16 different kidney cell-types using spatial transcriptomics. (A) H&E staining and DBSCANs of kidney sections from female KDM6A^Ctrl^ mice (n=4) and female KDM6A^TubKO^ mice (n=4). (B) Quality control metrics. (C) Aggregated UMAP showing 16 different kidney cell-types detected: ascending loop of Henle in cortex (ALOH), collecting duct (CD), connecting tubules and collecting ducts (CNT_CD), distal tubules and connecting tubules (DT_CNT), glomerular cells, immune cells, injured proximal tubule (InjPT), injured proximal tubule segment 3 cells (InjPT3), juxtaglomerular cells, loop of Henle in outer medulla (LOH (IOM)), proximal tubule S1 segment (PTS1), proximal tubule S1S2 (PTS1S2), proximal tubule S2 (PTS2), proximal tubule S3 (PTS3), vasa recta, and vasculature. (D) Annotated spatial clusters in kidney sections from a female KDM6A^Ctrl^ mouse and a female KDM6A^TubKO^ mouse. Note InjPT and InjPT3 cells were most abundant at the outer medulla in both KDM6A^Ctrl^ and KDM6A^TubKO^ kidneys, whereas tubule cell vacuolization was present at the outer cortex of female KDM6A^TubKO^ kidneys. Accordingly, InjPT and InjPT3 cells were considered likely to reflect tissue ischemia at the time of harvest rather than being reflective of the consequences of gene knockout.

Because the phenotype of female KDM6A^TubKO^ mice was characterized by glucosuria and decreased SGLT2 protein levels and SGLT2 is expressed in proximal tubule S1 and S2 segments, we focused our pseudobulk differential gene expression analysis on the comparison between female KDM6A^TubKO^ and KDM6A^Ctrl^ proximal tubule epithelial cells. Examination of PCA plots revealed clear genotype separation between female KDM6A^TubKO^ and KDM6A^Ctrl^ PTS1, PTS1S2, PTS2 and PTS3 cells (S. I. Fig. 4). Interestingly, pseudobulk differential gene expression analysis identified similar patterns of gene dysregulation across proximal tubule segments (Fig. 7A). For instance, *Elolv2*, which encodes Elongation of very long chain fatty acids protein 2 (ELOVL2), was differentially downregulated in female KDM6A^TubKO^ PTS1, PTS1S2, PTS2 and PTS3 cells (Fig. 7A). Other genes important for cellular metabolism, ion transport and/or antioxidant signaling also downregulated in multiple proximal tubule cell-types included: *Ccdc198*, *Afm*, *Clic6*, *Fam149a*, *Kidins220*, and *Apom* (Fig. 7A). These gene expression changes were associated with downregulation of pathways involved in amino acid, small molecule and glycoprotein metabolism, membrane transport and cellular respiration, along with induction of cellular activation and immune response pathways across proximal tubule cells (Fig. 7B).

**Figure 7.**
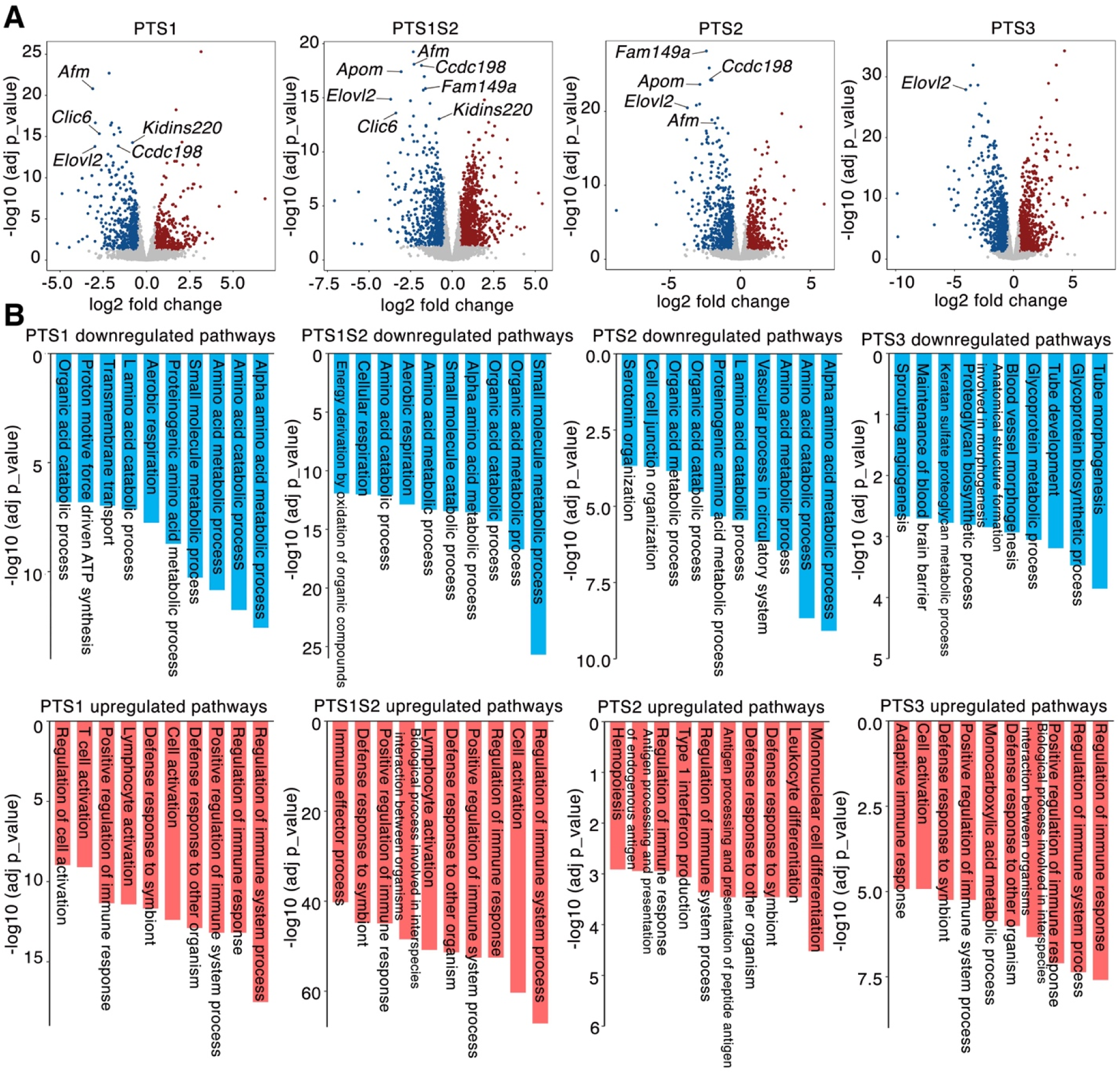
Knockout of KDM6A from tubule cells in female kidneys causes downregulation of genes and pathways involved in cellular metabolism and ion transport, and upregulation of immune responses in proximal tubule cells. (A) Volcano plots showing differentially expressed genes between female KDM6A^TubKO^ (n=4) and KDM6A^Ctrl^ kidneys (n=4) in PTS1, PTS1S2, PTS2 and PTS3 cells. Red circles indicate upregulated genes and blue circles indicate downregulated genes, adjusted *P* value < 0.05, log_₂_ fold-change > 0.5. *Elovl2* was downregulated with KDM6A knockout in each cell-type, and *Ccdc198*, *Afm*, *Clic6*, *Kidins220*, and *Apom* were downregulated in more than one cell-type. (B) Gene set enrichment analysis (GSEA) showing the top 10 Gene Ontology (GO) terms for upregulated and downregulated pathways in the comparison of female KDM6A^TubKO^ and KDM6A^Ctrl^ PTS1, PTS1S2, PTS2 and PTS3 cells.

### Absence of KDM6A in female mice causes ultrastructural and metabolomic changes suggestive of disrupted proximal tubule mitochondrial and metabolic dysfunction

Given that absence of KDM6A in female mice resulted in disruption of homeostatic transcriptional programs in tubule epithelial cells, we next examined proximal tubule ultrastructure by transmission electron microscopy. This survey revealed the presence of vacuoles and heterogeneous electron-dense lysosomes in female KDM6A^TubKO^ proximal tubule cells (Fig. 8A), with occasional areas of cytoplasmic disarray and mixed content vesicles suggestive of blocked cargo trafficking (Fig. 8B); with occasional degenerated cells with dense aggregates (Fig. 8B). By morphometry, mitochondria of female KDM6A^TubKO^ proximal tubule cells demonstrated a significant, albeit modest, diminution in elongation (increased roundness (Fig. 8C and decreased aspect ratio (Fig. 8D)) and increased circularity (Fig. 8E) suggestive of mitochondrial stress.

**Figure 8.**
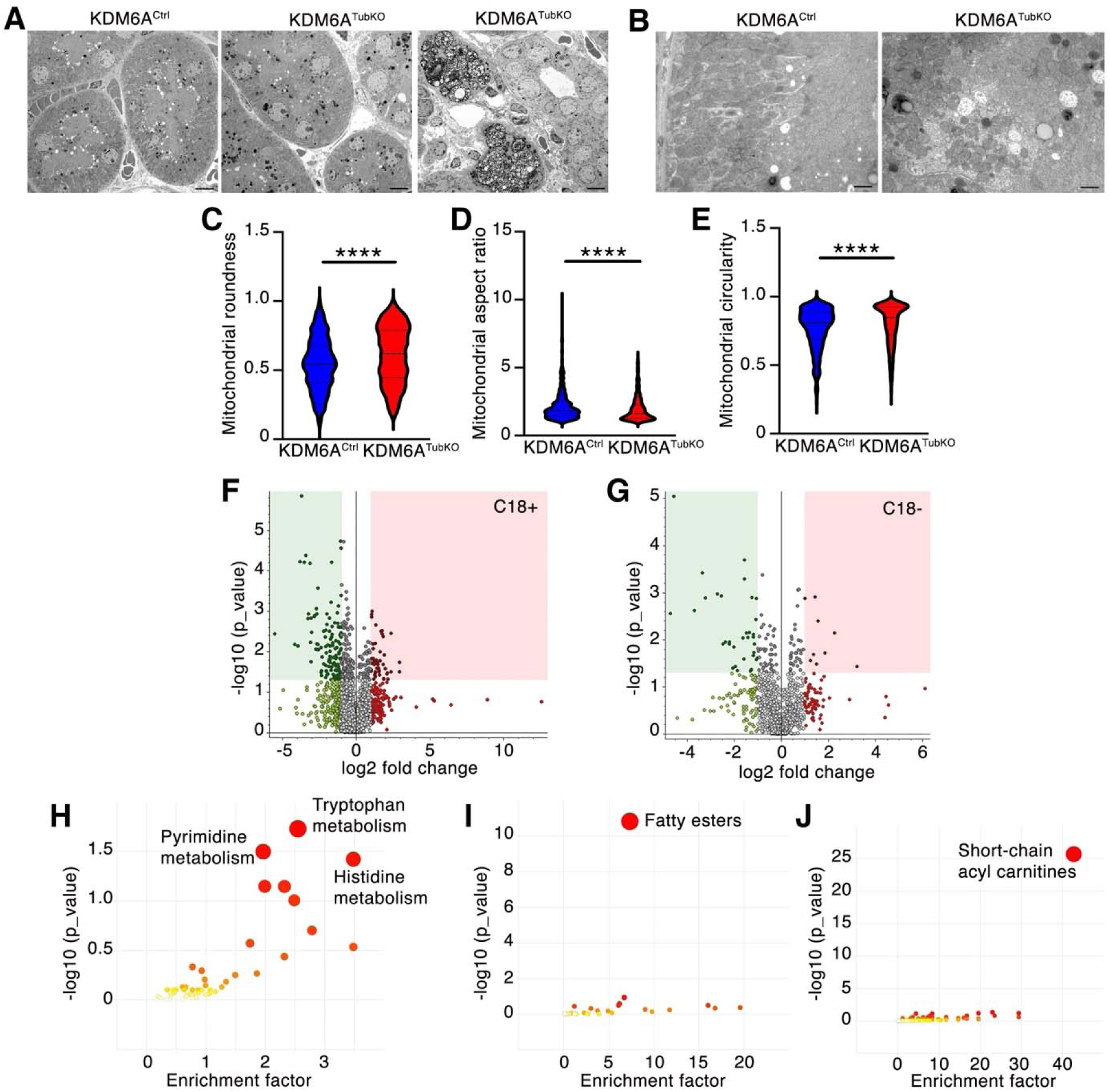
Knockout of KDM6A causes tubule cell vacuolization, mitochondrial rounding and metabolic dysfunction in female mice. (A and B) Representative transmission electron micrographs showing accumulation of vacuoles and electron dense bodies and increased mitochondrial circularity in proximal tubule cells of female KDM6A^TubKO^ mice; with focal areas of marked cellular degeneration (right). n=4/group. (A) scale bar = 5 µm. (B) scale bar = 1 µm. (C-E) Roundness (C), aspect ratio (D) and circularity (E) measured in 573 mitochondrial cross sections in proximal tubule cells from KDM6A^Ctrl^ mice and 688 mitochondrial cross sections in proximal tubule cells from KDM6A^TubKO^ mice. (F and G) Volcano plots showing differentially altered metabolites in the kidneys of female KDM6A^TubKO^ mice in comparison to KDM6A^Ctrl^ (F, C18+; G, C18-). n=4/group. (H-J) KEGG pathway analysis of C18+ metabolites showing pathway enrichment for all metabolites (H), lipid class (I) and lipid subclass (J). Values are mean ± S.D. \**P* < 0.05 by two-tailed Student *t* test.

Lastly, we compared metabolites of the kidneys of female KDM6A^TubKO^ and KDM6A^Ctrl^ mice using untargeted metabolomics. In total we detected 2252 compounds in positive mode (477 compounds identified) and 1089 compounds in negative mode (212 compounds identified). Examination of principal component analysis (PCA) plots revealed group separation of metabolites in positive ionization mode (S. I. Fig. 5); with volcano plot analysis demonstrating that the majority of differentially abundant metabolites identified in positive ionization mode were reduced in abundance in KDM6A^TubKO^ kidneys (Figs. 8F and G). The decreased metabolites included those linked to tryptophan, pyrimidine, and histidine metabolism (Fig. 8H), with lipidomic profiling revealing decreased abundance of fatty esters (Fig. 8I) and short-chain acylcarnitines (Fig. 8J) consistent with altered fatty acid utilization and impaired mitochondrial substrate metabolism. These metabolomic shifts align with spatial transcriptomic evidence of decreased amino acid metabolic and cellular respiration programs in proximal tubule cells, indicating the occurrence of proximal tubule metabolic dysfunction when KDM6A is absent in females.

## DISCUSSION

Here we report transcriptional and metabolic reprogramming, cellular injury and spontaneous glucosuria in female mice when KDM6A is deleted from tubule epithelial cells. When the kidney is stressed by occlusion of the ureter exiting the contralateral kidney these alterations manifest as a Fanconi renotubular syndrome-like picture. Mechanistically, KDM6A maintains homeostatic genes and pathways in proximal tubule cells including those involved in cellular metabolism, ion transport, respiration and antioxidant signaling; whereas its absence provokes an inflammatory response. Males, on the other hand, express the KDM6A paralog UTY and are largely protected from the deleterious consequences of KDM6A deletion. The findings provide molecular level insights into sex-dependent differences in kidney (patho)physiology, and they define KDM6A as an essential regulator of tubule cell homeostasis and stress resilience in females.

Our discovery arose from an experiment we conducted in follow up to our prior observation of mostly inconsequential effects of deletion of KDM6A from tubule epithelial cells in male mice either normally or when the kidney is stressed by UUO.^16^ In our subsequent experiments we made several new discoveries. Firstly, fibrosis of obstructed kidneys is augmented by KDM6A absence from tubule cells in female mice but not males. Secondly, female but not male KDM6A^TubKO^ mice develop glucosuria, tubule epithelial vacuolopathy and SGLT2 downregulation. Thirdly, tubule cell-specific KDM6A knockout in female mice with UUO causes polyuria, along with *Tnf* induction and interstitial fibrosis in the unobstructed contralateral kidney with heightened urinary loss of phosphate and other electrolytes. Fourthly, knockout of KDM6A from tubule epithelial cells of female mice induces a molecular signature indicative of metabolic dysfunction and disrupted tubule cell homeostasis.

From a non-reductionist stance, we considered it unlikely that the role of KDM6A in preserving tubule cell integrity be limited to any single signaling pathway. Accordingly, we used spatial transcriptomics to gain insights into the coordinated transcriptional consequences of KDM6A deletion, focusing on these consequences in proximal tubule cells. We observed downregulation of several homeostatic genes. For instance, *Elovl2* was downregulated across proximal tubule cell-types. *Elovl*2 is a master controller of lipid metabolism and its epigenetic regulation is highly associated with aging.^35^ *Ccdc198* and *Afm* expression levels were diminished in PTS1, PTS1S2 and PTS2 KDM6A^TubKO^ cells. *Ccdc198* encodes Coiled-Coil Domain Containing 198 (CCDC198), also called Factor Associated with Metabolism and Energy (FAME) and has recently been linked to kidney homeostasis,^36^ whereas *Afm* encodes the vitamin E-binding protein afamin and lower serum afamin levels are associated with kidney function decline.^37^ *Clic6* and *Kidins220* levels were also lower in KDM6A^TubKO^ PTS1 and PTS1S2 cells. *Clic6* encodes the redox-sensitive anion channel Chloride Intracellular Channel 6 (CLIC6) and *Kidins220* polymorphisms are associated with endstage kidney disease and obesity.^38^ *Fam149a and Apom* expression were diminished in PTS1S2 and PTS2 cells. FAM149A is a positive regulator of Nuclear factor erythroid 2-related factor 2 (Nrf2) antioxidant signaling.^39^ Apolipoprotein M (the protein encoded by *Apom*) functions as a carrier for sphingosine-1-phosphate and is linked to protection against acute kidney injury (AKI) and chronic kidney disease (CKD).^40^ Downregulation of these, and other homeostatic genes and pathways, was accompanied by diminished amino acid metabolism, increased rounding of proximal tubule mitochondria (a morphological hallmark of mitochondrial stress),^41^ induction of inflammatory genes and pathways and reduced pathways supporting nutrient utilization, mitochondrial activity and cellular homeostasis. Integration of these observations that span from transcriptional and ultrastructural changes in single cells, through kidney metabolism and tissue structure, to kidney physiology underscores the pivotal importance of KDM6A for the maintenance of female kidney health.

Situating our findings in the context of the extant literature, Han et al recently reported that knockout of KDM6A from tubule cells causes salt-sensitive hypertension in mice, including in males.^42, 43^ However, the authors of those reports do not comment on the presence or absence of glucosuria in the mice they studied. Of relevance, the investigators employed mice expressing Cre recombinase under the control of the *Ksp-cadherin* gene promoter to induce tubule cell gene deletion,^42^ and *Ksp-cadherin* has been reported to have little or no expression in the proximal tubule.^44, 45^ Thus, our discovery was only made possible by: i) the intentional comparison of male and female KDM6A^TubKO^ mice under stressed conditions; and ii) the use of *Pax8*-Cre^+^ mice to achieve tubule cell deletion, Pax8 being expressed throughout the kidney tubules, including in proximal tubule cells.^16^ The present findings also build on those of a recent study in which KDM6A inhibition counteracted the deleterious effects of high glucose on mitochondrial respiration in proximal tubule epithelial cells from males in vitro.^8^ These findings, together with our earlier report of altered peroxisome proliferator-activated receptor (PPAR) pathway signaling in the kidneys of male KDM6A^TubKO^ mice,^16^ indicate that KDM6A has some functional role in tubule epithelial cells in males. However, the magnitude of the phenotypic difference between male and female KDM6A^TubKO^ mice herein reported emphasizes that the actions of KDM6A are highly sex-dependent. This supposition is congruent with observations in embryology,^17, 18^ and with observations made in hematopoietic stem and progenitor cells (HSPC), where female HSPC KDM6A knockout mice develop myelodysplasia and splenic erythropoiesis, and male HSPC KDM6A knockout mice are phenotypically unremarkable.^46^ The findings are also reminiscent of those of a recent report that KDM6A knockout dysregulates differentiation programs in the epidermis, sebaceous glands and hair follicles in females but not males.^47^ In that report, the authors attributed the protected phenotype of males to the compensatory action of UTY.^47^

KDM6A was first recognized as a demethylase whose enzymatic activity is necessary for HOX gene cluster regulation during development.^10–12^ Unlike KDM6A, UTY has been suggested to have little, if any, demethylase activity.^17, 48^ For instance, human UTY may have weak demethylating activity ^48^ and mouse UTY has been reported to be catalytically inactive.^17^ However, KDM6A also has non-enzymatic actions, catalytically inactive KDM6A being sufficient for embryonic stem cell differentiation ^18^ and mouse viability.^49^ These non-enzymatic actions likely reflect the scaffolding function of KDM6A. For example, KDM6A is a member of the MLL3/MLL4 COMPASS complex that deposits H3K4me1 at enhancer regions and promotes p300-mediated H3K27 acetylation.^13–15^ UTY has also been reported to associate with COMPASS complexes,^17, 50^ although the literature implicating UTY as a member of COMPASS is currently far less than it is for KDM6A. Furthermore, both KDM6A and UTY have been reported to associate with several other proteins and chromatin-modifying complexes that regulate gene transcription, aside from roles either as H3K27me3 demethylases, or as members of the MLL3/MLL4 COMPASS complex.^9, 15, 17^ Recent studies comparing epigenomic changes caused by KDM6A and/or UTY knockout report genome-wide losses of H3K27 acetylation with minimal changes in H3K27 methylation.^47, 51^ Given these epigenomic changes, together with the emerging evidence of functional redundancy between KDM6A and UTY and the minimal demethylating actions of UTY, a picture is emerging that many of the biologically critical actions of KDM6A are likely to be independent of its catalytic activity.

Despite the strength of the phenotype of female KDM6A^TubKO^ mice, especially under conditions of added stress, the present study has limitations that warrant emphasis. We have reported for the first time that *Uty* is expressed in male tubule epithelial cells in vivo and it is upregulated in response to UUO, and we have proposed that UTY may provide at least partial compensation for KDM6A absence in male kidneys. However, defining the physiological actions of UTY in the kidney extends beyond the scope of the current work. Likewise, we have not untangled the relative contributions of genomic and endocrine (androgen) factors in protecting the kidneys of male mice from tubule cell KDM6A-specific knockout. Furthermore, whereas we observed no change in kidney H3K27me3 levels in female KDM6A^TubKO^ mice, we have not confirmed that the actions of KDM6A in preserving tubule epithelial cell homeostasis are independent of its demethylating activity. These limitations notwithstanding, the present report is timely and significant because, situated alongside our previous work and that of others,^8, 16, 42, 43, 47, 51, 52^ it firmly establishes KDM6A as a sex-dependent regulator of kidney tubule cell homeostasis and stress resilience.

In summary, KDM6A is a critical determinant of sex-dependent biological responses in the kidney. Knockout of KDM6A causes transcriptional reprogramming of tubule epithelial cells in females that manifests as glucosuria, vacuolopathy, and stress intolerance; whereas absence of KDM6A is relatively inconsequential in males. KDM6A/UTY are a dynamically regulated X-Y pair with at least partial functional overlap, likely now also extended to partial overlap of function in the kidney.

## ACKNOWLEDGEMENTS

The authors thank Dr. Laurette Geldenhuys (Department of Pathology, Dalhousie University, Halifax, Nova Scotia, Canada) for histological advice. Transmission electron microscopy was performed with support from Biotechnology and Biological Sciences Research Council (BBSRC) grant BB/R013942/1. Spatial transcriptomics was performed by the Princess Margaret Genomic Centre (RRID SCR_027524). Bioinformatics support was provided by the Canadian Center for Computational Genomics (C3G), a Genomics Technology Platform (GTP) supported by the Canadian Government through Genome Canada. The graphical abstract was generated with BioRender.com.

## DATA AVAILABILITY

Data are available from the corresponding author upon reasonable request. Spatial transcriptomics data have been deposited with GEO (accession number GSE336198). Metabolomic data have been deposited with Metabolomics Workbench, Study ID ST005038, DOI: http://dx.doi.org/10.21228/M8D86N.

## FUNDING

These studies were supported by project grants from the Canadian Institutes of Health Research (CIHR; PJT 166083, 183810 and 203724) and by a John R. Evans Leaders Fund Award from the Canada Foundation for Innovation (38214) to A.A.. L.Y.Q.H. was supported by a Yow Kam-Yuen Graduate Scholarship in Diabetes Research from the Banting and Best Diabetes Centre (BBDC), an Angels Den PhD Scholarship from St. Michael’s Foundation and a BBDC - Novo Nordisk Studentship. D.T.T. was supported by a Novo Nordisk - BBDC Postdoctoral Fellowship. M.Z.S. was supported by a Keenan Post-Doctoral Scholarship from St. Michael’s Foundation and is supported by a CIHR Fellowship. This work was supported by funding to C.H.B. for “The Metabolomics Innovation Centre” from Genome Canada through the Genomics Technology Platform (265MET and MC4T), and by grant #42495 to C.H.B. from the Canada Foundation for Innovation via the Major Sciences Initiatives Fund. Metabolomics Workbench is supported by NIH grant U2C-DK119886 and OT2-OD030544 grants. A.A. holds the Keenan Chair in Medicine from St. Michael’s Hospital and University of Toronto. Work in the Advani Lab is supported by the RDV Foundation and the Fenella Foundation.

## DISCLOSURES

D.A.Y. is a scientific cofounder and consultant for Fibrocor Therapeutics. A.A. has received research support through his institution from Boehringer Ingelheim. Other authors declare no competing interests.

## CONTRIBUTION STATEMENT

S.N.B. and A.A. conceived the study. S.N.B., D.A.Y. and A.A. designed the experiments.

L.Y.Q.H., S.N.B., D.T.T., M.Z.S., S.L.A. and Y.L. performed the experiments. L.Y.Q.H., A.P. and

A.A. analyzed the data. E.V.P. and C.H.B. performed the metabolomics. L.Y.Q.H. and A.A. drafted and revised the paper. All authors approved the final version of the manuscript.

## DECLARATION OF GENERATIVE AI AND AI-ASSISTED TECHNOLOGIES IN THE MANUSCRIPT PREPARATION PROCESS

During the preparation of this work, the authors used GEMINI and ChatGPT for some literature searches and some minor text editing. After using these tools, the authors reviewed and edited the content as needed and take full responsibility for the content of the published article.

## SUPPORTING INFORMATION FIGURE LEGENDS

**S. I. Figure 1.**
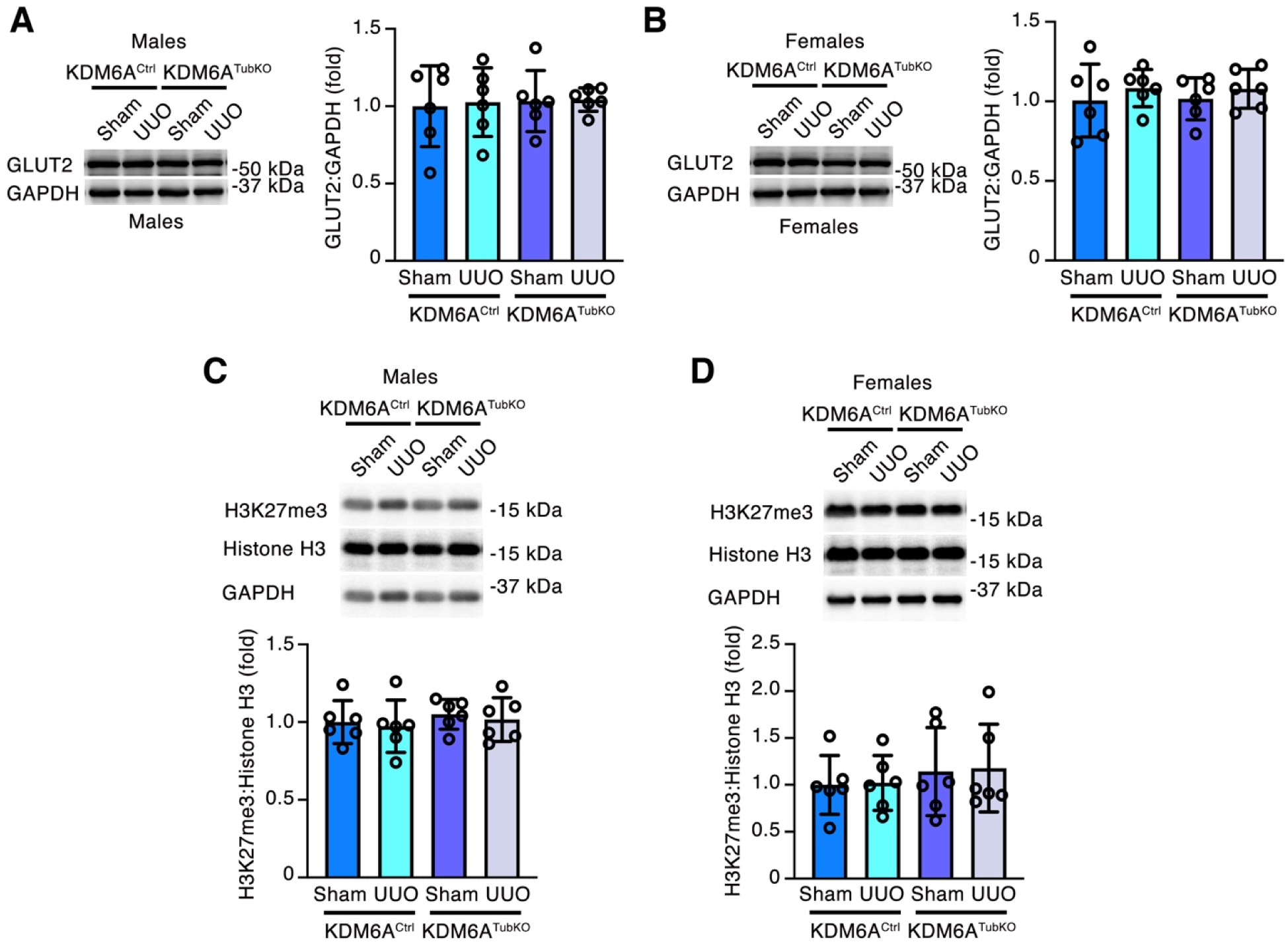
Immunoblotting for GLUT2 (A and B) and H3K27me3 (C and D) in the kidneys from male (A and C) and female (B and D) KDM6A^Ctrl^ and KDM6A^TubKO^ mice taken from sham-operated animals or the unobstructed (right) kidneys of mice 14 days after unilateral ureteral obstruction (UUO). n=6/group.

**S. I. Figure 2.**
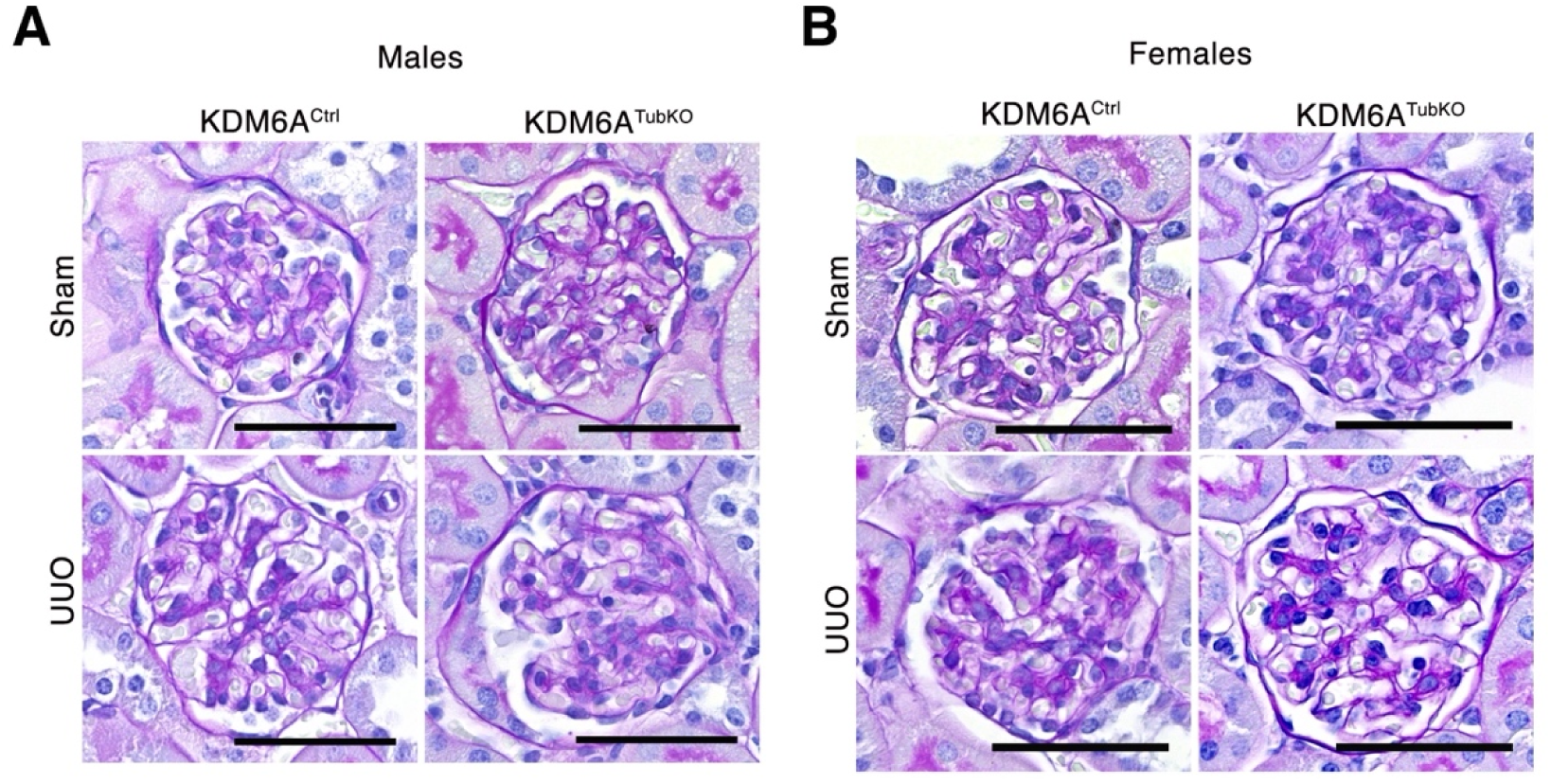
Representative Periodic acid-Schiff stained kidney sections from male (A) and female (B) KDM6A^Ctrl^ and KDM6A^TubKO^ mice taken from sham-operated animals or the unobstructed (right) kidneys of mice 14 days after unilateral ureteral obstruction (UUO). n=4/group. Original magnification x 400. Scale bar = 50 µm.

**S. I. Figure 3.**
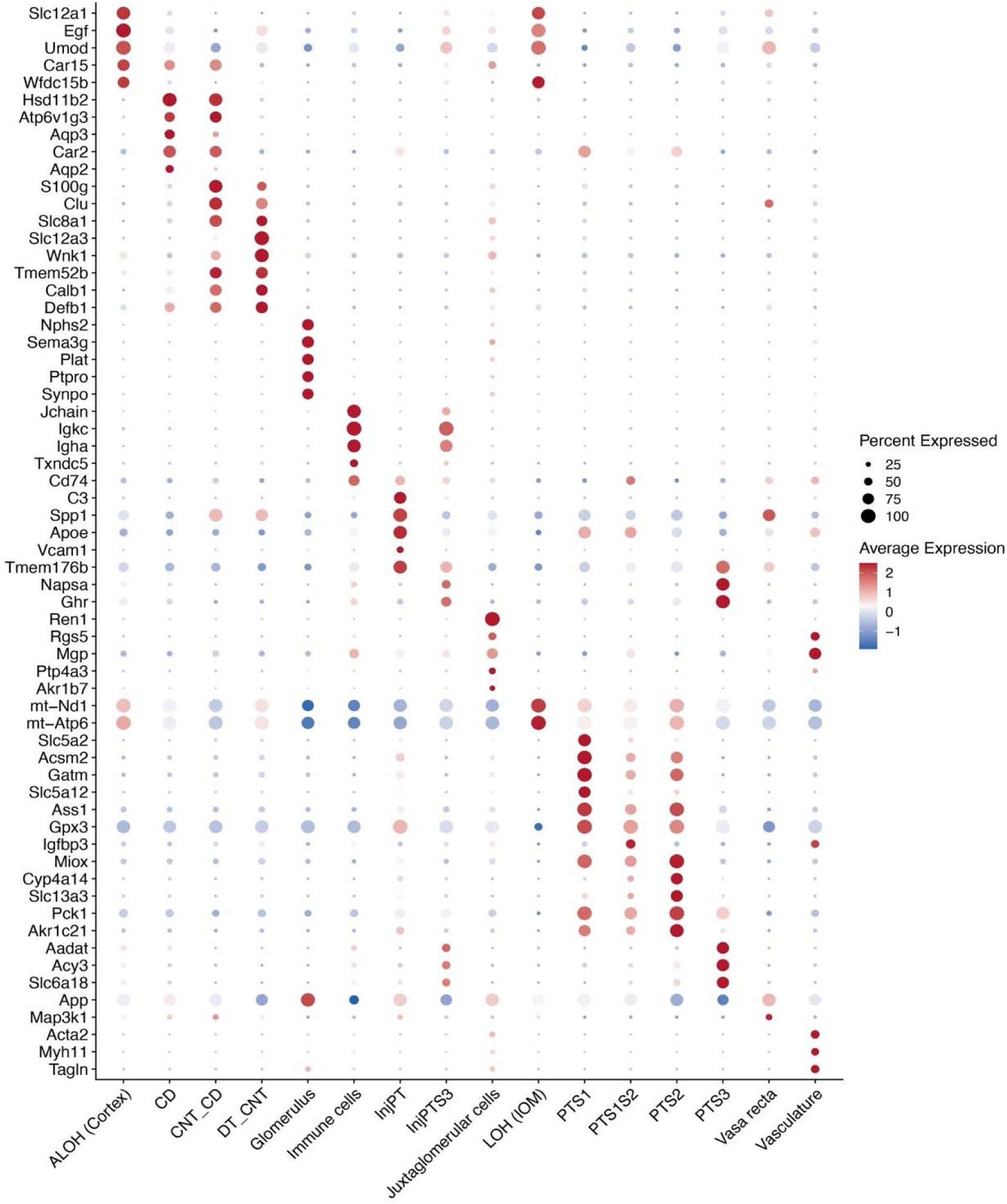
Marker gene expression for the 16 cell-types identified with 10x Genomics Visium HD spatial transcriptomics.

**S. I. Figure 4.**
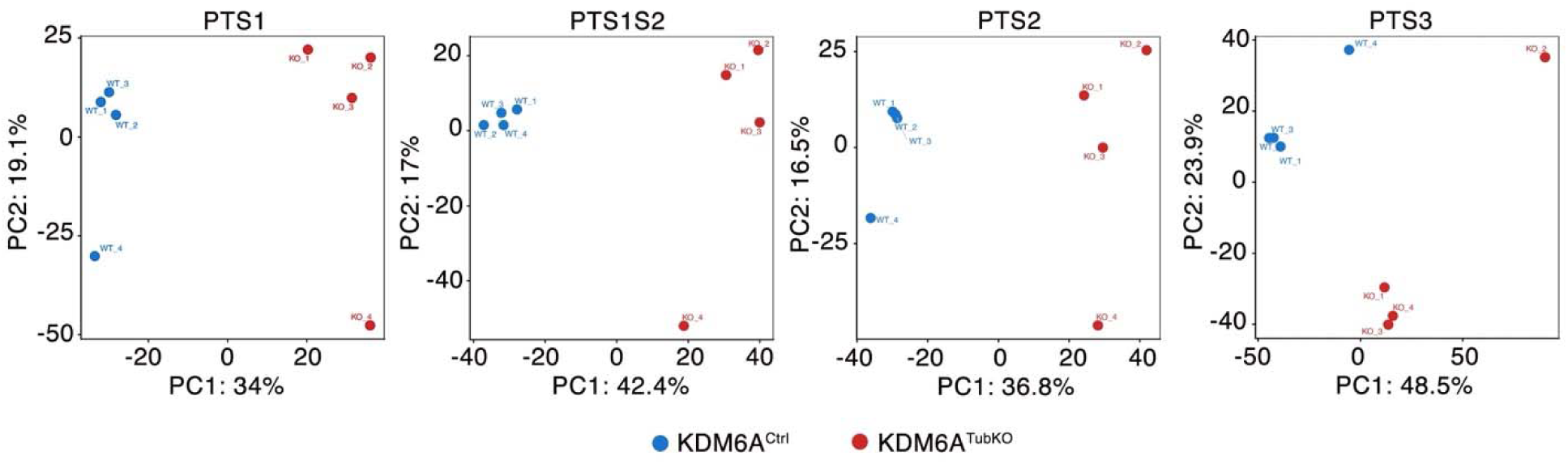
Principal component analysis (PCA) plots for PTS1, PTS1S2, PTS2 and PTS3 cells in female KDM6A^Ctrl^ (WT) and KDM6A^TubKO^ (KO) kidney sections using 10x Genomics Visium HD spatial transcriptomics.

**S. I. Figure 5.**
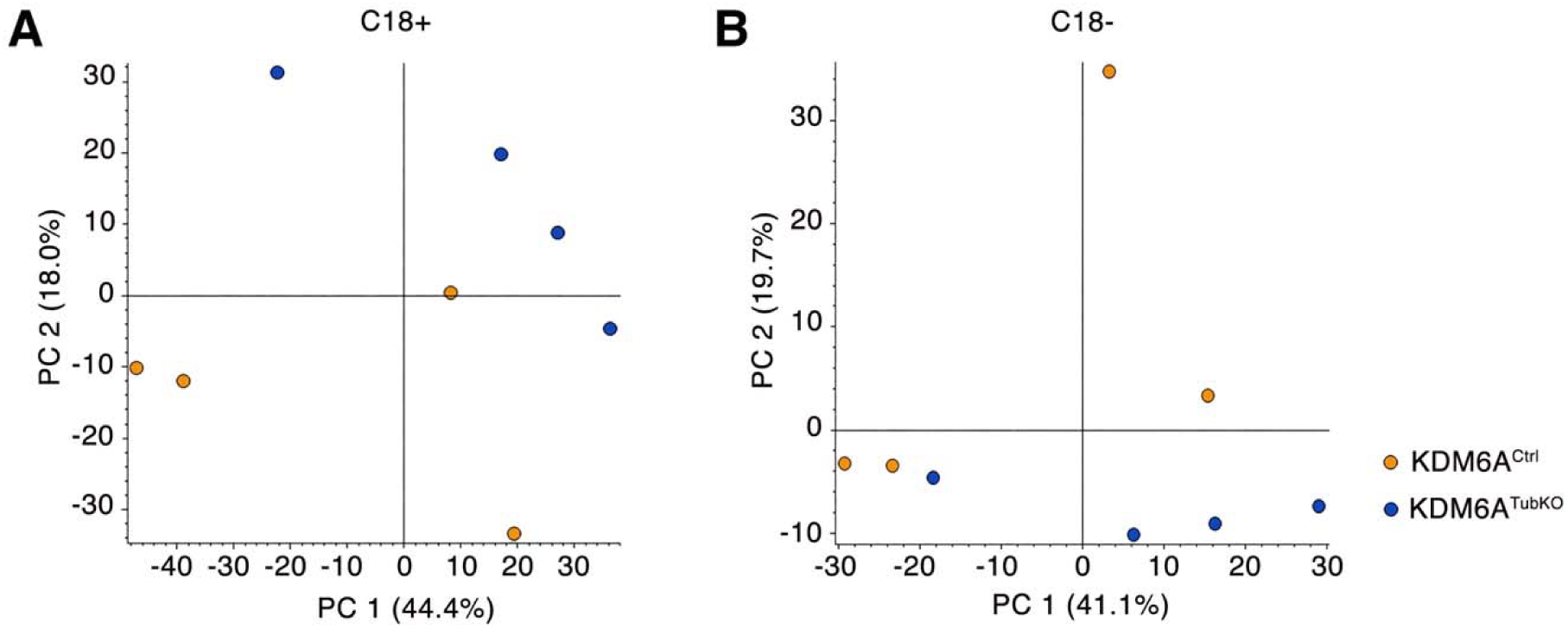
Principal component analysis (PCA) plots for female KDM6A^Ctrl^ and KDM6A^TubKO^ kidney sections using untargeted metabolomics.

**S. I. Table 1.**
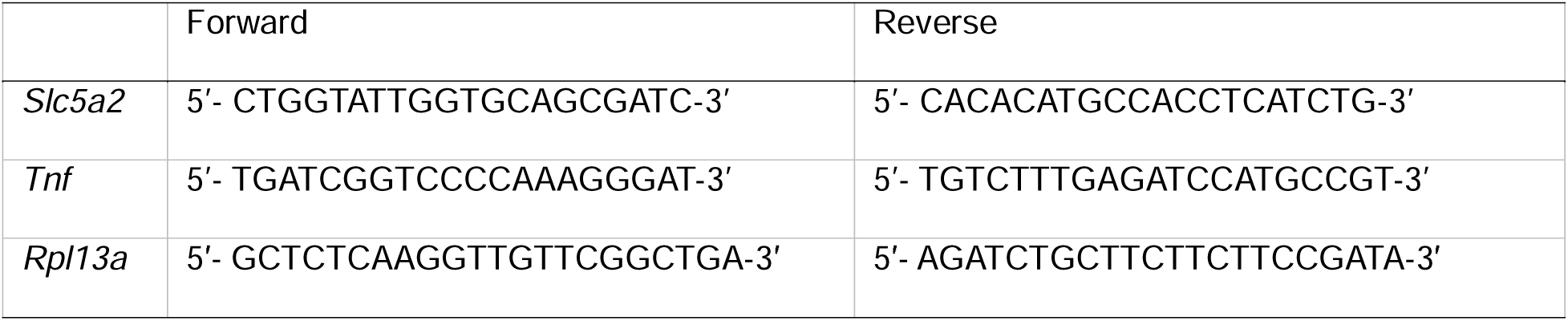
Primer sequences used for qRT-PCR.

**S. I. Table 2.** Quality control metrics for spatial transcriptomics of kidney sections from female KDM6A^Ctrl^ and KDM6A^TubKO^ mice.

|  | KDM6A<sup>Ctrl</sup> (n=4) | KDM6A<sup>TubKO</sup> (n=4) |
| --- | --- | --- |
| Number of 8 µm binned squares under tissue | 472,245 | 395,890 |
| Mean reads per 8 µm bin | 1789.2 | 1677.7 |
| Mean UMIs per 8 µm bin | 328.8 | 366.2 |
| Total genes detected | 17,858 | 17,690 |
| Reads mapped to probe set | 98.9% | 99.0% |
| Reads mapped confidently to probe set | 98.7% | 98.8% |
| Reads mapped confidently to the filtered probe set | 98.5% | 98.6% |
| Reads half-mapped to probe set | 0.2% | 0.2% |
| Reads split-mapped to probe set | 0.0% | 0.0% |
| Number of reads | 844,934,726 | 664,173,556 |
| Valid barcodes | 86.6% | 88.4% |
| Valid UMI sequences | 99.9% | 99.9% |
| Sequencing saturation | 77.5% | 73.7% |
| Q30 bases in barcode | 97.4% | 97.5% |
| Q30 bases in probe read | 97.0% | 97.1% |
| Q30 bases in UMI | 96.4% | 96.3% |
| Fraction of bins under tissue 8 µm | 67.2% | 56.4% |
| UMIs per sq mm of tissue | 5,137,056 | 5,721,765 |
| Reads per sq mm of tissue | 24,042,699 | 22,998,624 |
| Fraction reads in squares under tissue | 99.3% | 99.3% |
| Number of cells | 152,408 | 152,823 |
| Reads in cells | 70.0% | 80.0% |
| UMIs in cells | 80.7% | 87.8% |
| Mean reads per cell | 3,873.2 | 3,374.5 |
| Median genes per cell | 540.0 | 562.0 |
| Median UMIs per cell | 694.0 | 723.0 |
| Median cell area (µm<sup>2</sup>) | 128.0 | 116.0 |
| Median nucleus area (µm<sup>2</sup>) | 28.0 | 28.0 |
| Maximum nucleus diameter (pixels) | 256 | 256 |

**S. I. Table 3.** Cell counts before and after quality control for spatial transcriptomics of kidney sections from female KDM6A^Ctrl^ and KDM6A^TubKO^ mice.

|  | Before_QC | After_QC |
| --- | --- | --- |
| KDM6B<sup>Ctrl</sup> 1 | 29,609 | 26,405 |
| KDM6B<sup>Ctrl</sup> 2 | 58,403 | 41,016 |
| KDM6B<sup>Ctrl</sup> 3 | 52,571 | 38,322 |
| KDM6B<sup>Ctrl</sup> 4 | 11,645 | 9,131 |
| KDM6B<sup>TubKO</sup> 1 | 52,490 | 43,595 |
| KDM6B<sup>TubKO</sup> 2 | 36,578 | 29,302 |
| KDM6B<sup>TubKO</sup> 3 | 42,389 | 30,514 |
| KDM6B<sup>TubKO</sup> 4 | 21,366 | 12,912 |

